# TLR4 signaling and macrophage inflammatory responses are dampened by GIV/Girdin

**DOI:** 10.1101/2020.08.29.273516

**Authors:** Lee Swanson, Gajanan D. Katkar, Julian Tam, Rama F. Pranadinata, Yogitha Chareddy, Jane Coates, Mahitha Shree Amandachar, Vanessa Castillo, Joshua Olson, Victor Nizet, Irina Kufareva, Soumita Das, Pradipta Ghosh

## Abstract

Sensing of pathogens by Toll-like receptor 4 (TLR4) induces an inflammatory response; controlled responses confer immunity but uncontrolled responses cause harm. Here we define how a multi-modular scaffold, GIV (a.k.a Girdin) titrates such inflammatory response in macrophages. Upon challenge with either live microbes or microbe-derived lipopolysaccharides (LPS, a ligand for TLR4), macrophages with GIV mount a more tolerant (hypo-reactive) transcriptional response and suppress pro-inflammatory cytokines and signaling pathways (i.e., NFkB and CREB) downstream of TLR4 compared to their GIV-depleted counterparts. Myeloid-specific gene depletion studies confirmed that the presence of GIV ameliorates DSS-induced colitis and sepsis-induced death. The anti-inflammatory actions of GIV are mediated *via* its C-terminally located TIR-like BB-loop (TILL)-motif which binds the cytoplasmic TIR-modules of TLR4 in a manner that precludes receptor dimerization; the latter is a pre-requisite for pro-inflammatory signaling. Binding of GIV’s TILL motif to other TIR modules inhibits pro-inflammatory signaling *via* other TLRs, suggesting a convergent paradigm for fine-tuning macrophage inflammatory responses.

**Significance:** To ensure immunity, and yet limit pathology, inflammatory responses must be confined within the proverbial ‘*Goldilocks* zone’. TLR4 is the prototypical sensor that orchestrates inflammatory responses through a series of well characterized downstream cascades. How TLR4 signals are confined remain incompletely understood. Using trans-scale approaches ranging from disease modeling in live animals, through cell-based interventional studies, to structure-guided biochemical studies that offer an atomic-level resolution, this study unravels the existence of a ‘brake’ within the TLR4 signaling cascade, i.e., GIV; the latter is a prototypical member of an emerging class of scaffold proteins. By showing that GIV uses conserved mechanisms to impact multi-TLR signaling, this work unravels a multi-scale point of convergence of immune signaling of broader impact beyond TLR4.

## Introduction

Macrophages are sentinel cells of the innate immune system; their location varies from peripheral blood to various organs including lungs, liver, brain, kidneys, skin, testes, and vascular endothelium. Consequently, dysregulated activation of macrophages impacts the outcome of diverse organ systems in a multitude of diseases (1).

Of the signaling pathways that modulate macrophage function, Toll-like receptors (TLRs) constitute a key signaling system; they recognize a wide variety of pathogen associated molecular patterns (PAMPS) and initiate acute inflammation through the production of inflammatory cytokines (2). Specialized TLRs for each class of PAMP allow fine tuning of the inflammatory response for efficient removal of the pathogen. The prototypic member, TLR4, efficiently senses gram-negative bacterial infections through recognition of the bacterial membrane component, lipopolysachharides (LPS). Binding of LPS to TLR4 triggers signaling cascades (e.g., NFkB and MAPK) that culminate in the production of pro-inflammatory cytokines (TNFα, IL-1β, IL-6, IL-12) and type-I interferons required for propagation of the inflammatory response and ultimately pathogen destruction (3). Although many components of the TLR signaling pathway have been well characterized, regulatory mechanisms that intricately balance pro-inflammatory and anti-inflammatory responses remain incompletely understood.

In this work, we reveal an unexpected role of GIV, a multi-modular scaffold protein and the prototypical member of the non-receptor <u>G</u>uanine nucleotide <u>E</u>xchange <u>M</u>odulator (GEM) family of proteins (4), as a key determinant of macrophage polarization and inflammatory cytokine expression. Because GIV binds and modulates G protein activity downstream of a diverse variety of ligand-activated receptors, e.g., growth factor and integrins [reviewed in (5, 6)], here we studied if and how it may impact the LPS/TLR4 signaling in the most relevant cell line, i.e., macrophages. We dissect the relevance of those findings in murine disease models and its broader relevance among other TLRs.

## Results and Discussion

### GIV is preferentially expressed in the myeloid cells of our immune system

Using publicly available protein expression databases (The Human Protein Atlas) we noted that GIV is highly expressed in several immune tissues including lymph nodes, appendix, spleen, bone marrow, and tonsil (**Supplementary Figure 1A**). An analysis of RNA-seq datasets curated by the NIH/NIAID-supported Immunological Genome Project (immgen.org) further confirmed that GIV (gene; CCDC88A) is most highly expressed in macrophages and dendritic cells, moderately expressed in B-cells and natural killer (NK) cells, and least expressed in T-cells (**Figure 1A**).

**Figure 1:**
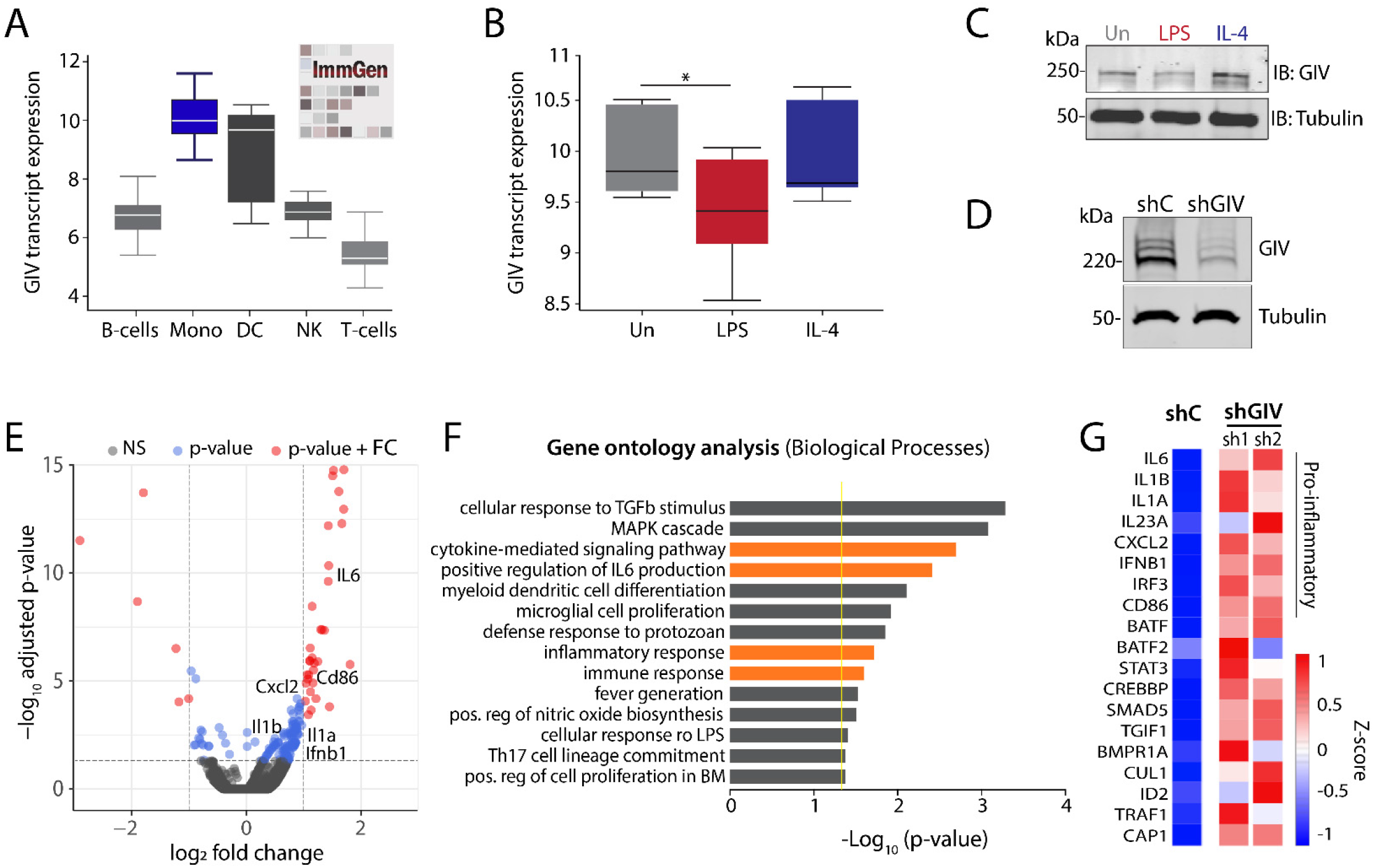
GIV/Gidrin expression is associated with pro-inflammatory gene programs in macrophages. **(A)** Box plots representing GIV (CCDC88A) transcript level in various immune cell populations. (Mono=monocyte, DC=dendritic cell, NK=natural killer cell) **(B)** Box plots representing GIV transcript levels in polarized CD14+ monocytes isolated from PBMCs (GSE35449) stimulated with either LPS (10 ng/ml) and IFNγ (200 U/ml) or IL-4 (1000 U/ml). **(C)** Immunoblot of GIV protein expression in polarized bone marrow-derived macrophages stimulated with either LPS (10 ng/ml) or IL-4 (20 ng/ml) for 24 hrs. **(D)** Immunoblot of RAW 264.7 macrophages depleted of GIV using shRNA. **(E)** Volcano plot of significantly (red) upregulated and downregulated gene transcripts in GIV-depleted (shRNA) RAW 264.7 macrophages stimulated with LPS (100 ng/ml, 6 hr) compared to controls (scrambled shRNA). Significance was determined using p-value < 0.05 and log2 fold change (FC) cutoffs. **(F)** Bar graph of significantly enriched biological processes determined by gene ontology (GO) analysis. Yellow line designates p=0.05 cutoff. Orange bars highlight biological processes relevant to macrophage inflammatory responses. **(G)** Heatmap of selected inflammatory gene transcript expression in GIV-depleted RAW 264.7 macrophages stimulated with LPS (100 ng/ml for 6 hr). shC=scrambled shRNA control; sh1 and sh2=two different GIV-targeting shRNA.

To explore possible functions of GIV in the immune cell type where it is most highly expressed, i.e., macrophages, we asked how its expression changes during macrophage polarization, which has classically been studied using a simplified nomenclature of reactive (a.k.a. M1, pro-inflammatory) *vs*. tolerant M2 (anti-inflammatory/healing). While the reactive state is important for engulfing and clearing invading pathogens and damaged cells and for mounting tailored inflammatory responses, the tolerant state is critical for restoring tissue homeostasis (7). An analysis of numerous human and mouse RNA-seq datasets (8) revealed that GIV expression is significantly decreased in LPS-stimulated (a widely used approach to induce M1 polarization (9)) but not IL-4-stimulated (a widely used approach to induce M2 polarization (10)) macrophages compared to controls (**Figure 1B, Supplementary Figure 1B, C**). These findings were validated in both murine bone marrow-derived macrophages (BMDMs) and RAW 264.7 macrophages stimulated either with LPS or IL-4 and subsequently assessed for GIV expression by immunoblotting (**Figure 1C, Supplementary Figure 1D**). We conclude that GIV is expressed in tissues with immune function and that high expression is seen in macrophages. In addition, GIV’s expression changes during macrophage polarization, i.e., it is suppressed in reactive macrophages but not in the tolerant ones.

### GIV dampens macrophage reactivity to live microbes and LPS

To study the role of GIV in macrophage inflammatory responses, we generated two model systems-- (i) a GIV-depleted RAW 264.7 macrophage cell line using short-hairpin RNA (shRNA) (**Figure 1D**) and (ii) a myeloid-specific conditional GIV knockout mouse, generated by crossing previously generated Girdin *floxed* mice (11) to LysMcre mice (**Supplementary Figure 2**). We asked if the presence or absence of GIV impacts macrophage responses and began by seeking insights from the transcriptome. GIV-depleted and control RAW 264.7 macrophages were stimulated with LPS and the relative levels of transcript expression were analyzed by RNA sequencing. We found that 150 genes were significantly upregulated, and 26 genes were significantly downregulated in GIV depleted macrophages compared to controls (**Figure 1E**). Gene ontology analysis performed using DAVID GO (https://david.ncifcrf.gov/) on the set of upregulated genes revealed 29 significantly enriched biological processes, whereas downregulated genes did not show such enrichment (**Figure 1F**). Most enriched pathways were involved in pro-inflammatory signaling and cytokine responses, including upregulation of IL-6, IL-1b, IL-1a, IL-23a, IL-17A, IL-12A, CXCL2, and IFNb1 (**Figure 1G**). These findings were confirmed by quantitative PCR and ELISA studies using GIV-depleted RAW 264.7 macrophages and peritoneal macrophages harvested from GIV KO mice (LysMcre) (**Figure 2A-D**). Increased expression of pro-inflammatory cytokines was consistently observed in both RAW 264.7 cells and peritoneal macrophages depleted of GIV (**Figure 2A-B**). Identical findings were observed in assays where we replaced LPS with the live microbes, *Escherichia coli* K12 strain and *Salmonella enteritica* serovar Typhimurium; both microbes induced a higher pro-inflammatory response in GIV-depleted macrophages compared to controls (**Figure 3A-C, Supplementary Figure 3C**).

**Figure 2:**
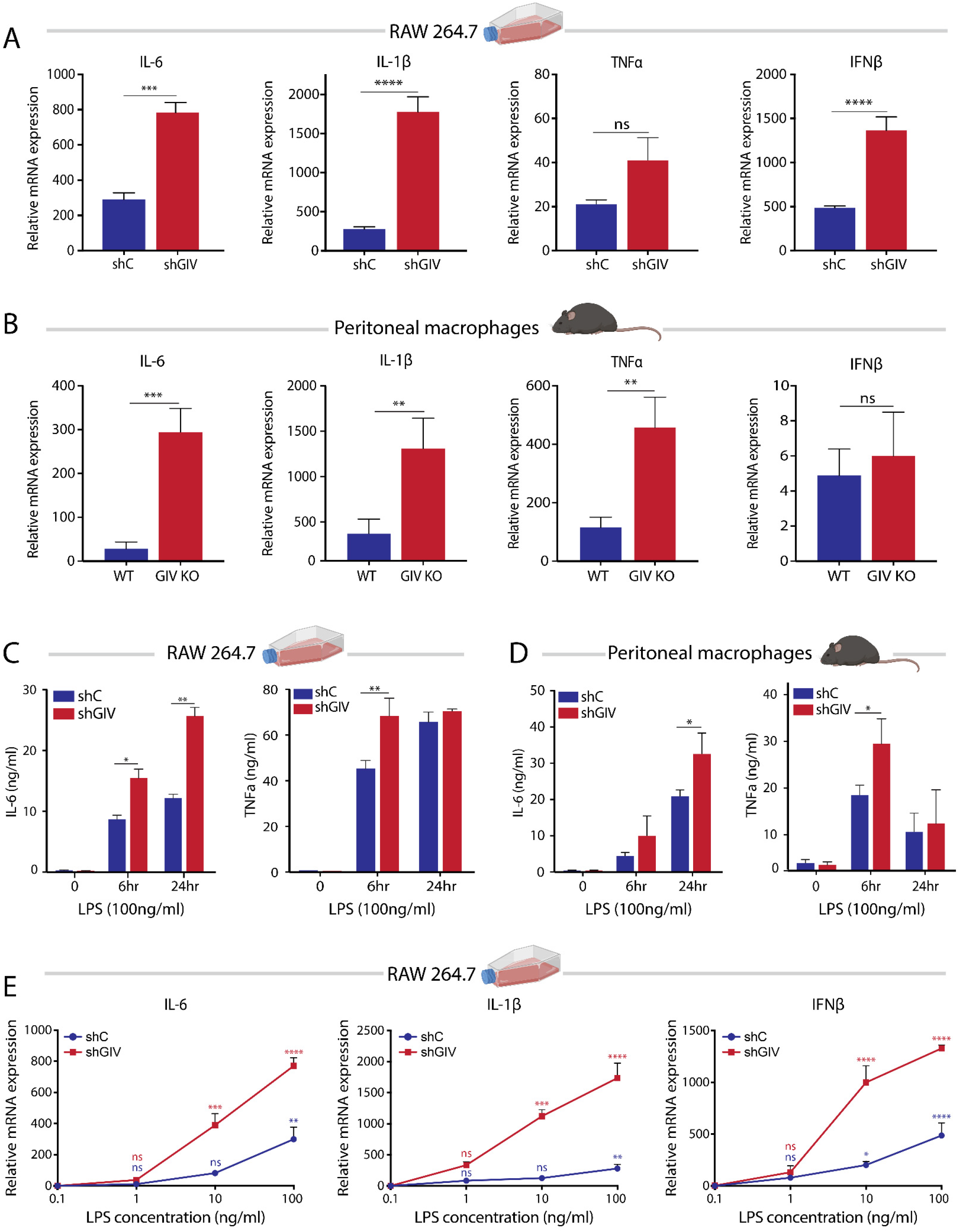
GIV depletion increases the magnitude and sensitivity of cytokine responses to LPS. **(A-B)** Bar graphs displaying cytokine transcript levels (qPCR) in GIV-depleted RAW 264.7 macrophages **(A)** or GIV KO peritoneal macrophages **(B)** stimulated with LPS (100ng/ml, 6hr) compared controls. **(C-D)** Bar graphs showing levels of secreted pro-inflammatory cytokines (ELISA) in GIV-depleted RAW 264.7 **(C)** or GIV KO peritoneal **(D)** macrophages stimulated with LPS. **(E)** Line graphs comparing sensitivity of cytokine transcript response to increasing doses of LPS (6 hr stimulation) in GIV-depleted RAW 264.7 macrophages compared to controls. All qPCR and ELISA results are from 3 independent experiments and displayed as mean ± S.E.M. Students t-test was used for two-parameter statistical analysis (A-B) and two-way ANOVA using Sidak’s multiple comparisons test was used for multi-parameter statistical analysis (C-E). (*;p ≤ 0.05, **;p ≤ 0.01, ***;p ≤ 0.001, ****;p ≤ 0.0001).

**Figure 3:**
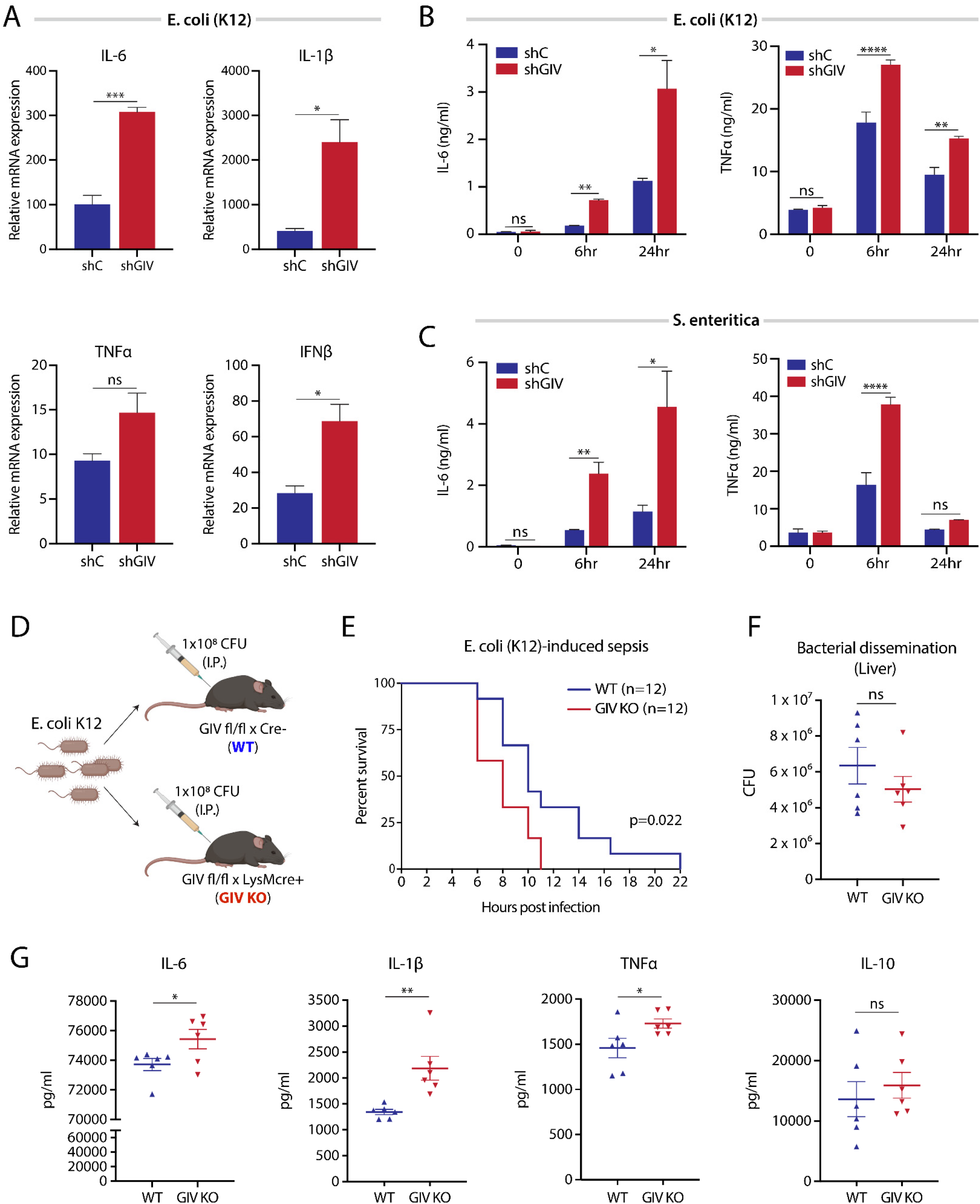
GIV depletion enhances pro-inflammatory cytokine response during live-microbe infection. **(A)** Bar graphs displaying cytokine transcript levels (qPCR) in GIV-depleted RAW 264.7 macrophages infected with live *E. coli* K12 (MOI=1) for 6hrs compared to controls. **(B-C)** Bar graphs showing levels of secreted pro-inflammatory cytokines (ELISA) in GIV-depleted or control RAW 264.7 macrophages infected with either *E. coli* K12 (MOI=1) or *S. enteritica* (MOI=10) for 6hrs. **(D)** Schematic of sepsis induced death mouse model. **(E)** Survival curve of GIV KO or WT mice infected with *E. coli* K1 strain RS218 (i.p.). Values expressed as percent survival. **(F)** Scatter plot of bacterial dissemination to liver following *E. coli* infection. **(G)** Scatter plots showing serum cytokine levels following *E. coli* infection. All qPCR and ELISA results are from at least 3 independent experiments and displayed as mean ± S.E.M.. Students t-test was used for two-parameter statistical analysis (A-B, F-G) and two-way ANOVA using Sidak’s multiple comparisons test was used for multi-parameter statistical analysis (B-C). (*;p ≤ 0.05, **;p ≤ 0.01, ***;p ≤ 0.001, ****;p ≤ 0.0001).

Intriguingly, compared to controls, the GIV-depleted RAW 264.7 and GIV KO peritoneal macrophages showed a reduction in the mRNA for IL-10 (**Supplementary Figure 3A**), a potent anti-inflammatory cytokine that promotes healing (12, 13). Thus, what emerged as a consistent finding is that compared to macrophages without GIV, those with GIV selectively suppress the pro-inflammatory responses to both forms of infectious stimuli, LPS and live microbes.

GIV may inhibit macrophage inflammatory responses either by reducing sensitivity or by inducing anergy (i.e., becoming refractive to repeated stimulation); to distinguish between the two, we carried out two commonly used assays, sensitivity at lower doses of LPS and anergy during repeated LPS challenges. Macrophages without GIV displayed increased reactivity to lower doses of LPS compared to controls (**Figure 2E**), indicating that GIV reduces sensitivity (or increases tolerance) of macrophages to LPS. When exposed to repeat doses of LPS, both control and GIV-depleted macrophages displayed LPS-induced anergy; even though GIV-depleted macrophages mounted a higher response than control cells during both exposures, repeat exposures elicited a weaker response than the first exposure in both cells (**Supplementary Figure 3B).** These findings indicate that the presence or absence of GIV may not impact anergy.

Taken together, these findings demonstrate that the presence of GIV in macrophages suppresses the pro-inflammatory gene signature, the production of pro-inflammatory cytokines (but not anti-inflammatory cytokine, IL-10), and reduces sensitivity to LPS without inducing anergy. We conclude that the physiologic role of GIV is to dampen pro-inflammatory macrophage responses.

### GIV ameliorates inflammation in murine models of colitis and sepsis

To explore the consequences of GIV’s role in macrophage polarization *in vivo*, we first utilized a sepsis-induced death model. WT or myeloid-specific GIV KO mice were infected with a lethal dose of *E. coli* (1×10^8^ CFU/mouse) and monitored for survival, levels of serum cytokines, and bacterial dissemination to spleen and liver (**Figure 3D**). GIV-depleted mice succumbed to sepsis-induced death significantly faster than controls (**Figure 3E**), and produced more pro-inflammatory cytokines (TNFα, IL-6, IL1-β, but not the healing-associated IL-10 cytokine) despite comparable amounts of microbial dissemination to peripheral organs (**Figure 3F-G**). Findings demonstrate that two major parameters in sepsis, i.e., ‘cytokine storm’ and death are suppressed in the presence of GIV. That the microbial counts were comparable in both groups suggests that the protective effect of GIV on sepsis-induced mortality is likely to be due to its ability to suppress the cytokine storm and unlikely to be confounded by overt changes in bacterial replication and/or defective clearance.

Next we assessed the role of GIV in colitis. We chose this as a disease model because of two reasons: (i) the gut, and more specifically, the colon is the largest reservoir for LPS-producing gram-negative bacteria (14), and (ii) hyperreactive immune responses by macrophages to gut microbes have been implicated in the initiation and perpetuation of colitis-associated syndromes such as inflammatory bowel disease (IBD; Crohn’s disease and Ulcerative Colitis) (15–17). We used the dextran sodium sulfate (DSS) mouse model of colitis because prior studies using this model have documented the importance of macrophage polarization states in limiting disease severity (18, 19). Because our prior observations demonstrate that GIV dampens macrophage inflammatory responses, we hypothesized that depletion of GIV in macrophages might exacerbate DSS-induced colitis. Mice were treated with DSS and monitored for changes in weight, stool consistency, rectal bleeding, colon length, colon tissue destruction, and immune infiltrates (**Figure 4A**). We found that GIV KO mice had increased weight loss, fibrotic shortening of the colon and disease activity index (DAI) compared to WT controls (**Figure 4B-D**). Histomorphological analysis of colon tissue sections revealed increased destruction of crypt architecture and immune infiltrates in GIV KO mice compared to WT controls (**Figure 4E-F**). Findings demonstrate that all the major parameters of severity of colitis were suppressed in the presence of GIV.

**Figure 4:**
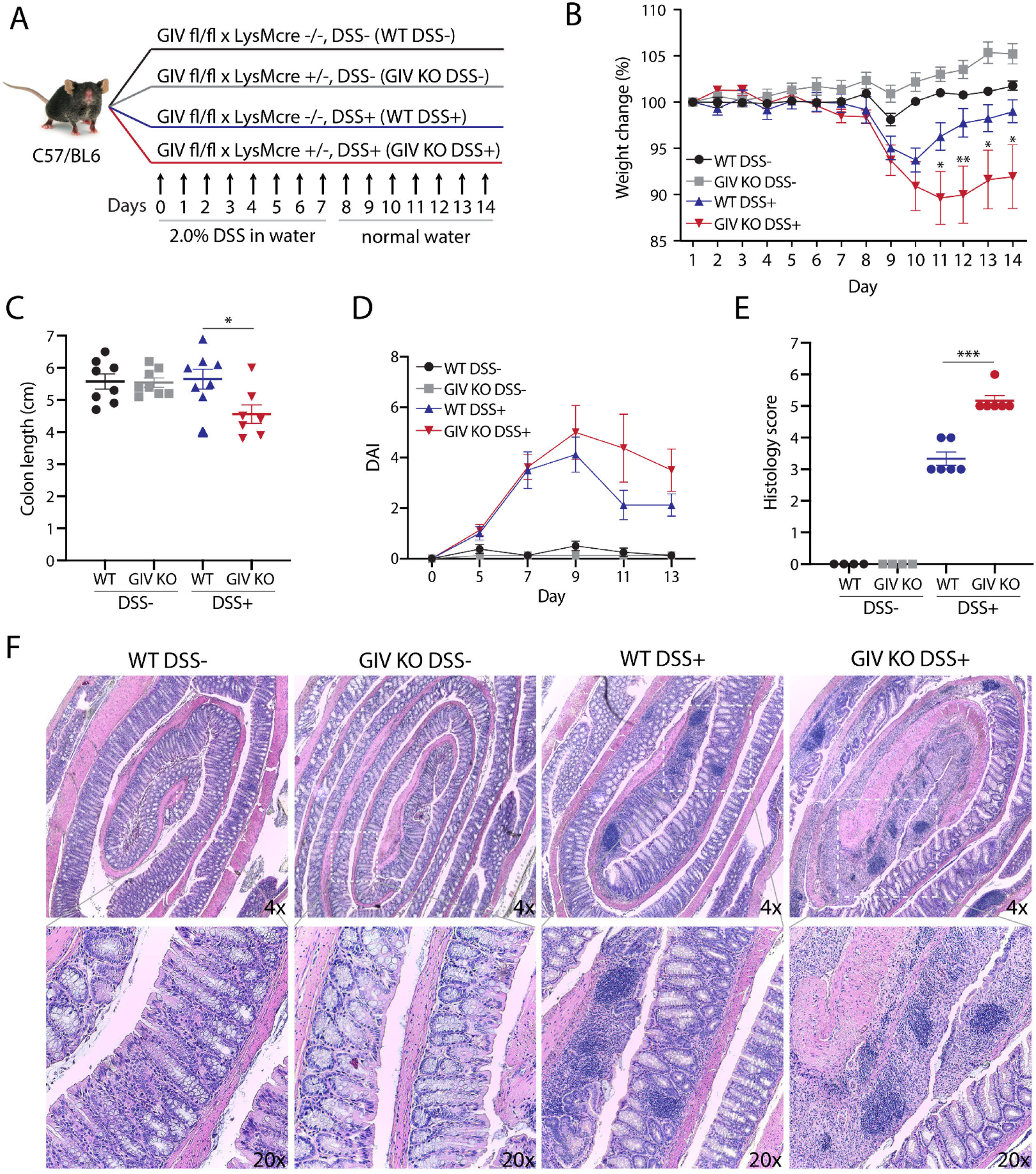
Myeloid cell specific GIV depletion exacerbates disease in DSS colitis: **(A)** Schematic outlining experimental design of DSS colitis model. **(B)** Line graph showing body weight change monitored daily during the course of acute DSS colitis. **(C)** Scatter plot of colon length assessed at d14 of DSS experiment. **(D)** Line graph of disease activity index (DAI) using stool consistency (0-4), rectal bleeding (0-4), and weight loss (0-4) as scoring criteria. **(E)** Scatter plot of histomorphological evaluation of inflammation by H&E stained colon tissues using inflammatory cell infiltrate (1–3), and epithelial architecture (1–3) as scoring criteria. **(F)** Representative images of colon tissue stained with H&E. Data displayed as mean ± S.E.M. and either one-way or two-way ANOVA using Tukey’s or Sidak’s multiple comparisons test was used to determine significance. (*; p ≤ 0.05, **; p ≤ 0.01, ***; p ≤ 0.001, ****; p≤ 0.0001).

These observations provide *in vivo* evidence for GIV’s role in restricting macrophage pro-inflammatory responses during microbial infection. Although the use of LysMcre for targeted depletion in macrophages is widely accepted, target protein depletion in other cell types including granulocytes, neutrophils, and dendritic cells cannot be ruled out (20); all of which are known to express GIV and play a role in the setting of sepsis (21) as well as in the pathogenesis of IBD (22, 23). However, taken together, the results from DSS colitis, acute sepsis, and *in vitro* cell stimulation assays (**Figures 1–4**) suggest that the phenotypes we observe, are at least in part due to GIV-depleted macrophages that are hyper-reactive.

### GIV suppresses LPS/TLR4-induced pro-inflammatory signaling pathways

Extensive work has gone into elucidating the signaling pathways downstream of LPS/TLR4 in macrophages(3, 24) (summarized in **Figure 5A**). GIV is known to modulate several of those pathways by linking G-protein signaling, *via* its GEM motif, to a multitude of cell surface receptors (reviewed in (25)). To investigate if GIV may also regulate TLR4 and/or the pro-inflammatory signaling pathways, control or GIV-depleted macrophages were stimulated with LPS and the signaling dynamics of key pathways (i.e., phosphorylation of NFkB, CREB, AKT, and MAPK) were assessed by immunoblot (**Figure 5B-C**). NFkB and CREB pathways, but not Akt, showed increased activation in GIV-depleted macrophages compared to controls, as examined by the ratio of phospho/total proteins. As for the MAPK pathway, GIV-depletion caused increased phosphorylation of p38 MAPK and increases in both phospho- and total ERK1/2 proteins. No appreciable difference was observed in JNK activation (**Supplementary Figure 4**). The pathways that were enhanced remained so even at 60 min, suggesting that GIV may inhibit negative feedback mechanisms required to dampen TLR4 signaling. We also observed enhanced CREB signaling in GIV-depleted macrophages as early as 5 min post stimulation, which is in agreement with the increase in sensitivity we observed in cytokine production **(Figure 2E)**. Because the expression of GIV did not decrease in control cells within 60 min of LPS stimulation [as it did after 24 hr stimulation **(Figure 1B, C)**], results indicate that GIV is required to suppress the pathways that were upregulated in GIV-depleted cells

**Figure 5:**
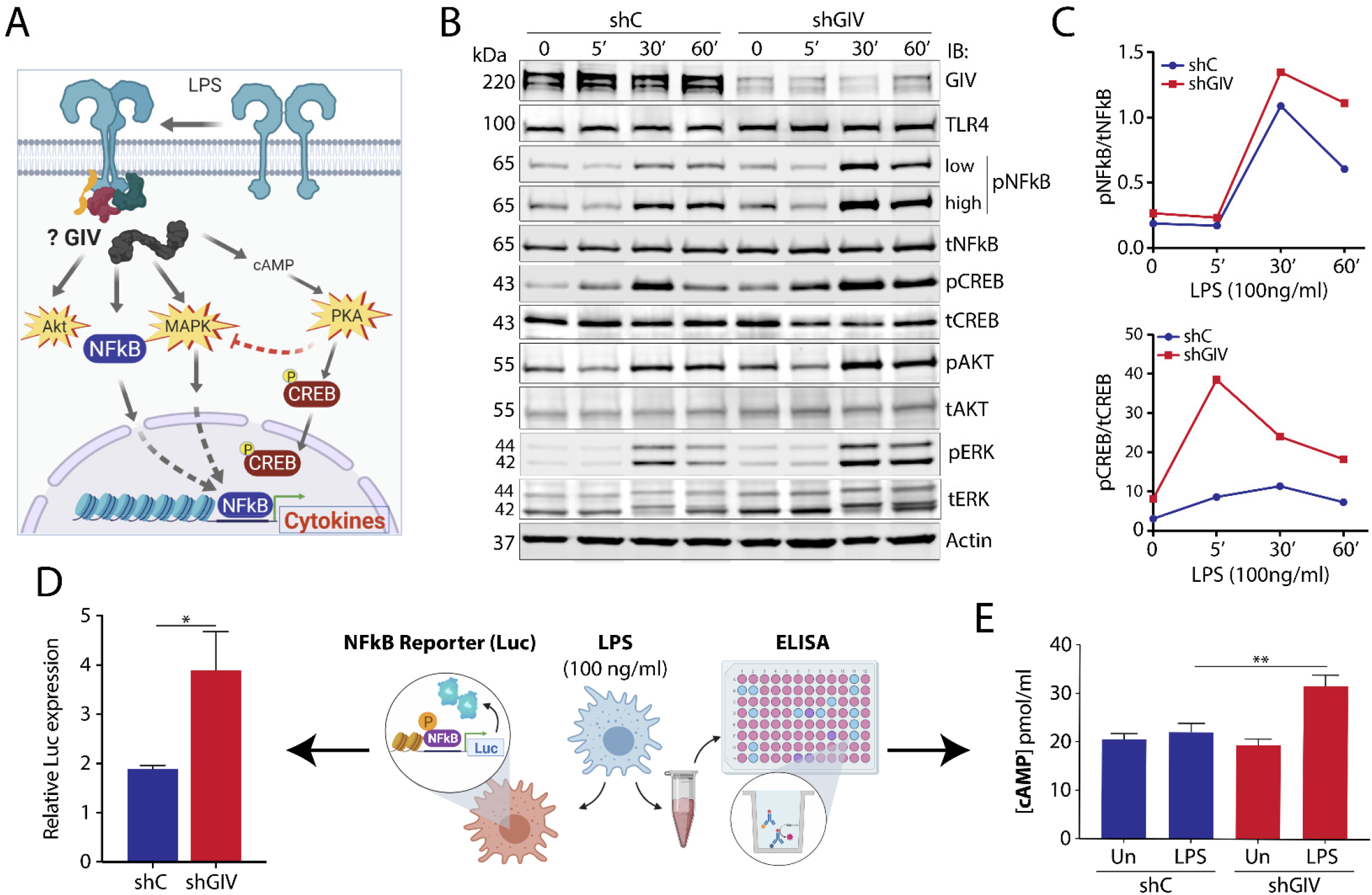
GIV depletion in macrophages enhances pro-inflammatory signaling pathways during LPS response. **(A)** Schematic highlighting GIV’s potential role in modulating signaling pathways downstream of TLR4. **(B)** Immunoblot of whole-cell lysates from GIV-depleted or control RAW 264.7 macrophages stimulated with LPS (100 ng/ml) and probed for activation of indicated signaling pathways. **(C)** Line graphs of representative densitometry values taken from signaling immunoblots. **(D)** Bar graph of relative NFkB activity in GIV-depleted or control RAW 264.7 macrophages stimulated with LPS (100 ng/ml, 1 hr) using NFkB luciferase reporter assay. **(E)** Bar graph of intracellular cAMP levels in LPS stimulated (100 ng/ml, 30 min) GIV-depleted or control RAW 264.7 macrophages. Results are from 3 independent experiments and displayed as mean ± S.E.M. Students t-test was used for two-parameter statistical analysis (D) and two-way ANOVA using Sidak’s multiple comparisons test was used for multi-parameter statistical analysis (E). (*;p ≤ 0.05, **;p ≤ 0.01).

We also found that GIV suppresses NFkB activity; upon LPS stimulation, NFkB activity was increased ~2-fold higher in GIV-depleted macrophages compared to controls, as determined by a well-established luciferase reporter assay (NanoLuc^®^ Promega) (**Figure 5D**). As for the observed increases in CREB phosphorylation, prior studies have implicated three parallel pathways downstream of TLR4 that are known to converge on CREB phosphorylation (26–29)-- (i) the cAMP→PKA pathway, (ii) the cAMP→Epac pathway, and (iii) the p38-MAPK cascade. Because GIV inhibits cAMP production by activating Gαi (30, 31), we hypothesized that cellular cAMP levels may increase during LPS stimulation in the absence of GIV. We found that cAMP levels were indeed elevated in GIV-depleted macrophages responding to LPS compared to controls (**Figure 5E**), indicating that GIV suppresses cellular cAMP in macrophages responding to LPS. We conclude that GIV-dependent suppression of cAMP downstream of TLR4 stimulation may represent a GPCR-independent pathway for modulation of cellular cAMP in macrophages responding to infections.

Taken together, these findings indicate that in macrophages responding to LPS, GIV specifically suppresses signaling within major pro-inflammatory signaling cascades, e.g., the NFkB→cytokine, cAMP→CREB and the MAPK/ERK pathways, but does not seem to significantly impact others, e.g., JNK and PI3K→Akt. Because the PI3K→Akt pathway promotes anti-inflammatory responses (32) and because JNK regulates macrophage development and survival (33), we conclude that GIV’s impact on TLR4 signaling is limited to those pathways that are directly related to cytokine production, but not cell fate.

### GIV directly binds the cytoplasmic TIR-module of TLR4, couples TLR4 to G protein pathways

To explore the mechanisms by which GIV modulates LPS/TLR4 signaling, and pinpoint where within the TLR4 signaling cascade GIV acts, we used a combination of biochemical assays. Published work from us and others has identified three critical features within the C-terminus of GIV: (i) a GEM motif that is required for interaction with and modulation of G proteins, (ii) distinct short linear interaction motifs (SLIMs) that couple GIV to receptor tyrosine kinases (RTKs), and Integrins (reviewed in (25)), and (iii) that all these SLIMs are packed within ~210 aa long GIV’s C-terminus, which is an intrinsically disordered protein (IDP) (34, 35) (**Supplementary Figure 5**). Because IDPs that fold/unfold thereby exposing/hiding SLIMs are known to impart plasticity to protein-protein interaction networks during signal transduction (36), we hypothesized that GIV may do something similar in TLR4 signaling. It may directly bind TLR4 through one SLIM, facilitate through another SLIM the assembly-disassembly of ternary receptor•GIV•Gαi complexes, and thereby dynamically shape post-receptor signaling. Immunoprecipitation of endogenous full length proteins from RAW 264.7 macrophages revealed that GIV forms a complex with TLR4 at steady-state (**Figure 6A**), and that such complexes were detected even ~30 min after LPS stimulation (**Supplementary Figure 7A)**. These findings show that the GIV•TLR4 interaction is constitutive, i.e., it is not significantly altered by ligand stimulation. *In vitro* pulldown assays using various fragments of recombinant His-GIV-CT and GST-tagged TLR4’s cytoplasmic – *Toll/interleukin-1 receptor* (*TIR*) module (TLR4-TIR; aa 676-835) proteins confirmed that the constitutive interaction between GIV and TLR4 observed in cells is direct. A ~110 aa stretch in GIV’s C-terminus (**Figure 6B**) and the cytoplasmic TIR module of TLR4 are sufficient for the interaction.

**Figure 6:**
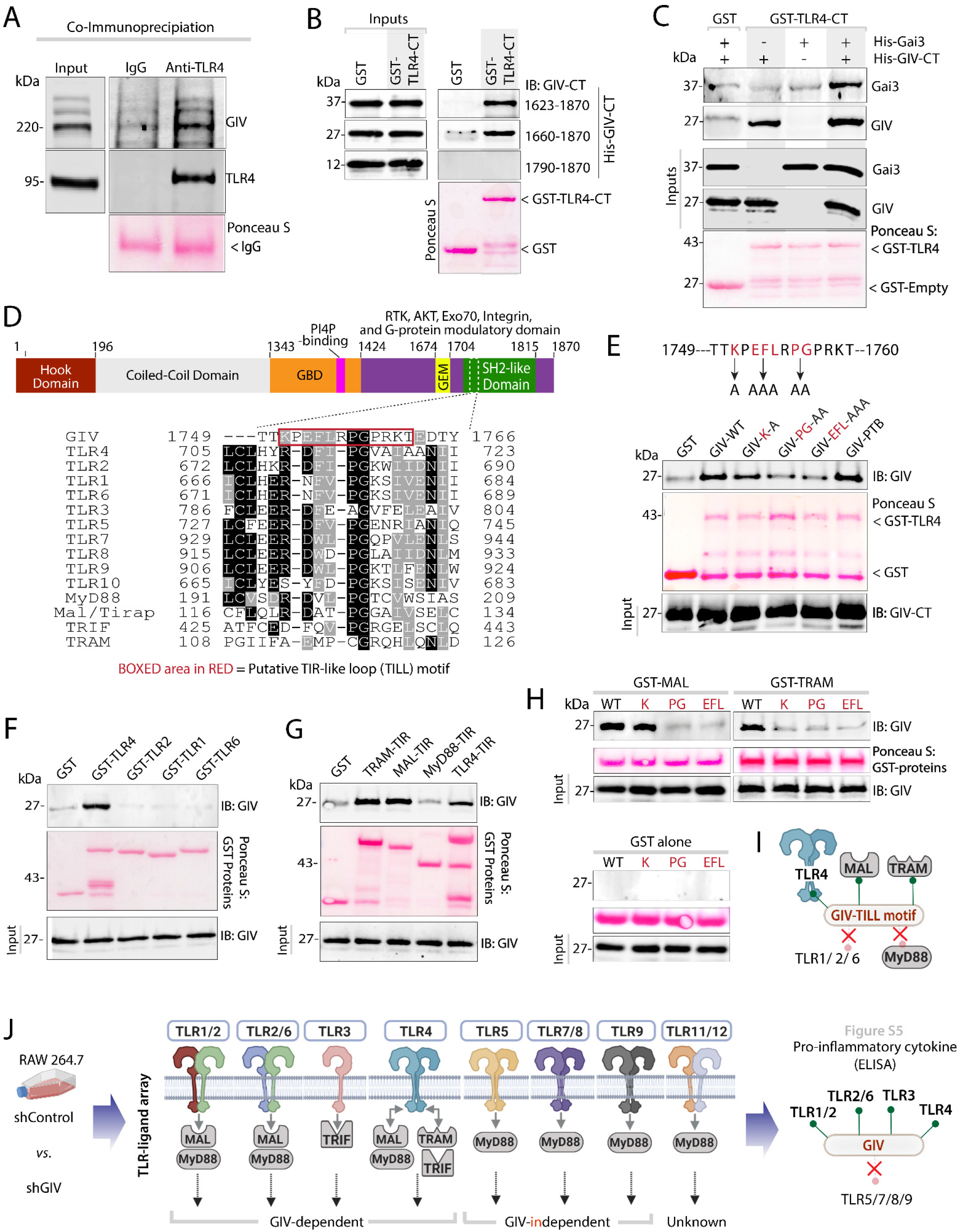
GIV directly interacts with TLR4 using a TIR-like loop (TILL) motif within its C-terminus and physically links TLR4 and Gai. **(A)** Endogenous TLR4 was immunoprecipitated from RAW 264.7 lysates. Bound complex of TLR4 and GIV were visualized by immunoblot. Equal loading of IgG and anti-TLR4 were confirmed by Ponceau S staining. **(B)** Various constructs of recombinant His-GIV-CT (~3mg) was used in GST pulldown assays with GST or GST-TLR4-TIR and bound GIV was visualized by immunoblot. **(C)** Recombinant His-GIV-CT (~3 mg) and His-Gai3 (~3 mg) were used in a GST pulldown assay with GST or GST-TLR4-TIR and bound GIV and Gai were visualized by immunoblot. **(D)** Sequence alignment showing short linear TIR-like loop (TILL) motif that is conserved between GIV and TIR containing proteins. **(E)** Recombinant His-GIV-CT or TILL mutants were used in a GST pulldown assay with GST or GST-TLR4-TIR and bound GIV was visualized by immunoblot. **(F-G)** Recombinant His-GIV-CT (~3mg) was used in a GST pulldown assay with various GST-TLR proteins **(F)**, GST-TIR adaptors **(G)**, and bound GIV was detected by immunoblot. **(H)** Recombinant His-GIV-CT TILL mutants were used in GST pulldown assays with TIR adaptor proteins and bound GIV was visualized by immunoblot. For all recombinant GST pulldown assays equal loading of GST proteins were confirmed by Ponceau S staining. **(I)** Schematic summarizing GIV•TIR interactions. **(J)** Schematic summarizing data from Supplementary Figure 6 investigating the impact of GIV on various TLR inflammatory responses.

Furthermore, using *in vitro* pulldown assay between recombinant TLR4-TIR (bait) and Gαi3 (prey) proteins in the presence or absence of GIV-CT, we confirmed that GIV facilitates the formation of a TLR4•GIV•Gαi ternary complexes (**Figure 6C**). Gαi3 bound TLR4 exclusively in the presence on GIV, suggesting that GIV acts as a physical link between TLR4 and Gαi, as it has been demonstrated to do for multiple RTKs (25) and for β1-integrin (35).

### A short linear motif (TILL) within GIV’s C-terminus binds multiple TIR-modules

Because TLR4 signaling relies on the ability of its cytoplasmic TIR module to assemble multimeric post-receptor homo- and heterotypic dimers via key contact sites, we hypothesized that the ~100 aa long stretch within GIV’s C-terminus may contain a SLIM that bears homology to one or more of such contact sites. Sequence alignment revealed a ~12 aa stretch with sequence homology to the BB-loop region of TIR domains (**Figure 6D**) which is also evolutionarily conserved (**Supplementary Figure 5**). The BB-loop is essential both for homo- and hetero-dimerization of TLRs and for the recruitment of TIR-domain-containing adaptors, and mutations in this loop inhibit TLR signaling (37). Additionally, a <u>T</u>IR-<u>l</u>ike BB <u>l</u>oop (henceforth, TILL) defined first in the cytoplasmic tail of IL17RA has been shown to be essential for NFkB and MAPK activation (38), adding support to the idea that TILL motifs can modulate inflammatory responses. To test if the newly identified putative TILL motif in GIV is required for binding TLR4, we replaced critical residues within GIV-TILL with Alanines (K^1749^A, EFL^1751-53^AAA, PG^1754-55^AA) and found that all three mutants showed decreased binding to TLR4. This motif appears to be specific because disruption of a neighboring SLIM that is ~13 aa upstream (PTB-binding motif, which enables binding to integrins) had no effect (35) (**Figure 6E**). These findings demonstrate that the direct interaction between GIV and TLR4-TIR is mediated via GIV’s C-terminally located TILL motif.

To assess what other TIR modules GIV may bind, we conducted GST pulldown assays with His-GIV-CT (prey) and GST-tagged TIR-modules found in the cytoplasmic tails of other TLRs (baits; TLR2, TLR1, TLR6) and in TLR-associated adaptor proteins (Mal, TRAM, TRIF, MyD88). Among the TLRs tested, TLR4 was the only one that could bind GIV (**Figure 6F**). GIV also bound the TIR adaptors MAL and TRAM, but not MyD88 (**Figure 6G**), indicating that GIV’s C-terminus can directly bind multiple TIR modules. Mutations within the TILL motif of GIV disrupted them all (**Figure 6H**), indicating that the TILL motif is necessary for these GIV•TIR interactions.

### GIV•TIR interactions aid in post-receptor signal integration, ligand specificity

Because GIV’s TILL motif bound multiple TIR-modules, but demonstrated selectivity by interacting with some (TLR4, MAL, TRAM), but not others (TLR1/2/3, MyD88) **(Figure 6I)**, we conclude that this pattern of binding may support two fundamental properties in immune response and signal transduction: (i) multi-TIR binding could mediate signal convergence/integration and receptor cross-talk to mount a consistent response regardless of the ligand, whereas (ii) selectivity between TLRs could impact ligand specificity. To study these two properties, we conducted a TLR-ligand screen where GIV-depleted or control RAW 264.7 macrophages were stimulated with ligands for various TLRs (**Figure 6J**). We observed a consistent increase in the production of pro-inflammatory cytokines in GIV-depleted macrophages stimulated with ligands for TLR1/2, TLRR2/6, and TLR3; however, the presence or absence of GIV did not impact responses to ligands for TLR5, TLR7/8, or TLR9 (**Figure 6J, Supplementary Figure 6)**, indicating that the latter are GIV-independent. We noted that the GIV-independent TLRs are capable of directly engaging MyD88, whereas those that are GIV-dependent engage with MAL, TRAM and TRIF. Because GIV binds MAL and TRAM, but not MyD88, these findings raise the possibility that GIV inhibits TLR signaling either by directly interacting with the cytoplasmic domain and preventing TIR-mediated receptor-dimerization or blocks the recruitment of other TIR-adaptors, or both. Alternatively, GIV could interact with TIR-adaptor proteins MAL and TRAM and sequester them from the cytoplasmic domain of other TLRs. Regardless of the exact mechanism(s) involved, it appears that GIV•TIR interactions aid in signal integration while maintaining specificity.

### Binding of GIV’s TILL motif to TLR4-TIR precludes TIR•TIR homotypic homodimerization, inhibits pro-inflammatory macrophage activation

We next sought a structural explanation of GIV’s binding to the TLR4 TIR domain and its effects on macrophage inflammatory signaling. Structural, biochemical, and computational studies have delineated two types of TIR•TIR assemblies–The so-called “homotypic” assembly involves inter-locking of the BB-loops regions of two TIR domains and is typically required for receptor homodimerization, whereas a “heterotypic” assembly occurs when the BB-loop of one TIR module binds the C-terminal helix of the other; such assembly is common in the process of recruitment of TIR-adaptors (39–41). Based on these observations, we hypothesized that the binding of GIV TILL motif to TLR4 may occur either at the BB-loop, mimicking and competitively disrupting homotypic TLR4-TIR dimers, or at the C-terminal helix, preventing the recruitment of TIR-domain containing adaptors. Approximate models of GIV TILL complexes with TLR4-TIR were built in both geometries (**Figure 7A; Supplementary Figure 7B**); interface residues were identified and compared across a panel of TIR domains that were shown experimentally to bind (or not) GIV-CT. The “homotypic” binding hypothesis provided a better explanation for the specificity of GIV’s interactions with TIR-proteins (**Figure 6F-H**): the alignment of interface residues from TLR4, MyD88, Mal, TRAM, TLR1, TLR2, and TLR6 revealed Q683, E685, and Y709 as contact sites specific to the GIV•TLR4 interface and closely conserved in other TIR domains that bind GIV but not in TIR domains that don’t (**Figure 7B-C**). By contrast, the heterotypic binding hypothesis (**Supplementary Figure 7B-C)** revealed that such binding could be accommodated but did not explain the binding specificity. In other words, while GIV’s TILL-motif sequence is compatible with both modes of binding, the homotypic, but not heterotypic, hypothesis is consistent with the observed selectivity of GIV for some TIRs (and not others). Model-guided mutagenesis confirmed that to be true, because replacement of contact site residues on the homotypic TLR4-dimer with corresponding residues found in TLR1/2/6 (i.e., TLRs that don’t bind GIV), reduced GIV’s ability to bind TLR4 (**Figure 7D**).

**Figure 7:**
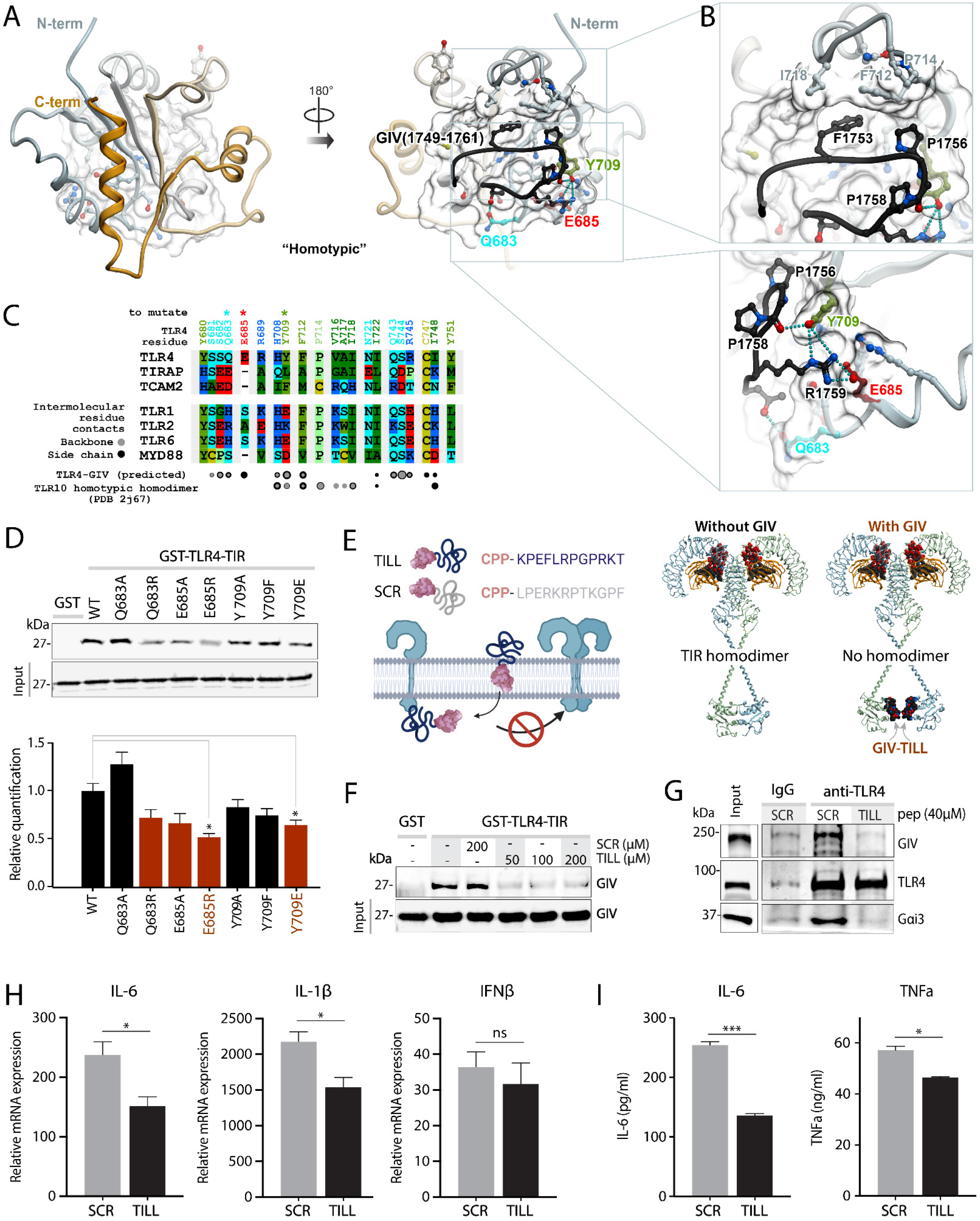
Identification of key contact sites between GIV’s TILL and BB-loop of TLR4. **(A)** A homology model of GIV’s TILL motif (black) bound to the BB-loop of human TLR4 (Grey) using a homotypic interface. **(B)** Magnified view of the predicted homotypic interface between GIV’s TILL and TLR4’s BB-Loop. **(C)** Sequence alignment of TLR4 with other TLRs and TIR-adaptors highlighting residues in GIV-TLR4 interaction interface that may confer binding specificity. **(D)** (Top) Recombinant His-GIV-CT was used in a GST pulldown assay with either WT GST-TLR4-TIR or GIV•TLR4 interface mutants and bound GIV was visualized by immunoblot. (Bottom) Bar graph of densitometry values from three independent experiments. **(E)** A model of TLR-bound GIV-TILL peptide showing the structural basis for obstruction of TIR•TIR homodimerization by GIV. **(F)** Schematic of cell-penetrating TILL peptide design. **(G)** Recombinant His-GIV-CT was used in GST pulldown assay with GST or GST-TLR4-TIR and increasing amounts of TILL-peptide or scrambled (SCR) control peptide. Bound GIV was visualized by immunoblot. Equal loading of GST proteins was confirmed by Ponceau S staining (Supplementary Figure 6). **(H)** Endogenous TLR4 was immunoprecipitated from RAW 264.7 lysates in the presence of either TILL-peptide or scrambled control. Bound complex of TLR4, GIV, and Gai were visualized by immunoblot. Equal loading of IgG and anti-TLR4 were confirmed by Ponceau S staining (Supplementary Figure 6). **(I-J)** Bar graphs displaying cytokine transcript levels (qPCR) (I) or secreted protein (ELISA) (J) in RAW 264.7 macrophages stimulated with LPS (100ng/ml, 6hr) in the presence of either TILL-peptide or scrambled control. Results are from 3 independent experiments and displayed as mean ± S.E.M. Students t-test was used to determine significance. (*;p ≤ 0.05, **;p ≤ 0.01, ***;p ≤ 0.001, ****;p ≤ 0.0001).

The homotypic model revealed key residues on GIV and TLR4 that facilitate binding: GIV’s K1750, P1756, and R1759 made contacts with TLR4’s Q683, E685, and Y709 respectively (**Figure 7B-C**). Identification of K1750 and P1756 as binding residues is supported by our biochemical assays (**Figure 6E**). It is also in keeping with prior literature supporting the essential role of the proline P1756 (which corresponds to proline 714 in TLR4 in humans) in the BB-loop of TLR4-TIR which is essential for TIR•TIR interactions (42). Tyrosine 709 is also known to form essential contacts in the TIR•TIR binding interface (43, 44); the latter opens the possibility that the GIV•TLR4 interaction may be phospho-regulated by tyrosine-based signals. The binding of GIV to TLR4 in a homotypic mode would preclude the assembly of homotypic homodimers of TLR4-TIR modules (**Figure 7E**). Because the latter is a pre-requisite for adaptor recruitment and for the initiation of pro-inflammatory signaling cascades (45), such binding is consistent with the experimental findings in this work.

To determine if the binding of GIV-TILL to TLR4 is sufficient to recapitulate the observed anti-inflammatory role of GIV in macrophages, we tested the ability of synthetic peptides mimicking this region to exogenously modulate macrophage inflammatory responses. Cell-penetrable peptides containing the TILL motif of GIV (KPEFLRPGPRKT) fused to the C-terminus of the Antennapedia peptide (RQIKIWFQNRRMKWKK (46); **Figure 7F**) were synthesized commercially. These peptides, representing the minimal required segment of GIV’s C-terminus were predicted to have two measurable consequences if they bound TLR4: (i) such binding should ‘displace’ GIV•Gαi complexes from TLR4, and (ii) binding should inhibit TLR activation and the production of pro-inflammatory cytokines. As expected, the GIV-TILL peptide, but not a control peptide with a scrambled sequence, but of same charge and residue composition, disrupted the interaction between His-GIV-CT and GST-TLR4-TIR in *in vitro* protein interaction assays (**Figure 7G**). The GIV-TILL peptide also displaced both full-length GIV and Gαi from immunoprecipitated TLR4 in RAW 264.7 macrophages (**Figure 7H**). When we investigated LPS-triggered inflammatory responses, macrophages incubated with GIV-TILL peptide had reduced expression of pro-inflammatory cytokines compared to control peptide with scrambled sequence (**Figure 7I-J**).

These results demonstrate that the GIV-TILL motif is both necessary and sufficient for binding to TLR4; such binding is sufficient to inhibit pro-inflammatory responses upon LPS stimulation in macrophages. These findings add support to the working model that GIV’s TILL motif binds and disrupts TLR4 signaling, thereby inhibiting pro-inflammatory cascades (see legend, **Figure 8)**.

**Figure 8:**
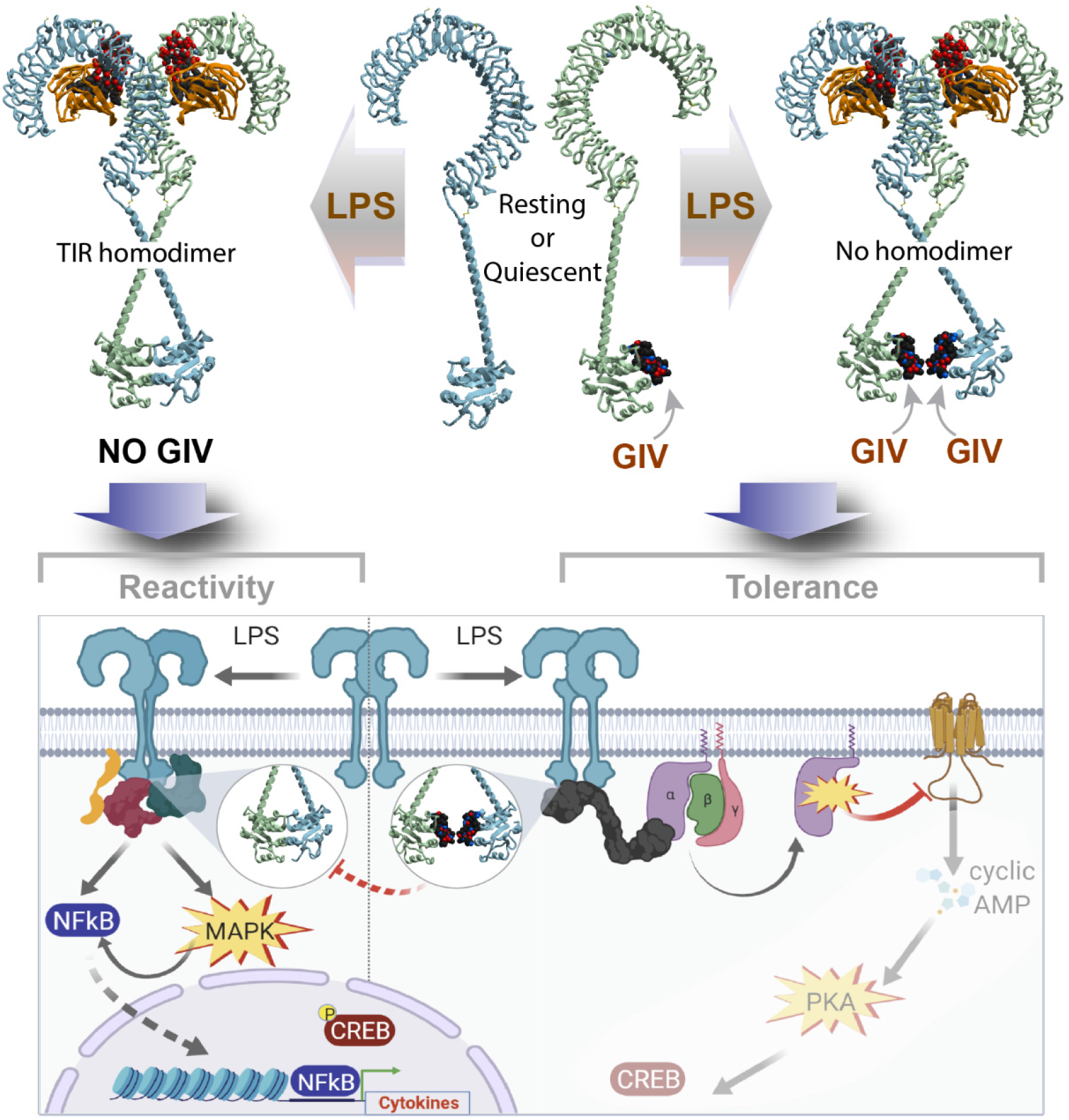
Summary of findings and working model. In macrophages expressing GIV (right), GIV constitutively and directly binds the cytoplasmic TIR domains of TLR4. Binding occurs under resting conditions (center, top), which reduces the sensitivity of macrophages to low concentrations of LPS stimuli. Binding of GIV to TLR4 also continues after ligand stimulation (right, top) and their mode of binding precludes receptor dimerization (via TIR•TIR homodimerization). These may serve as a potential mechanism for the inhibition of proinflammatory signaling cascades during an LPS/TLR4 inflammatory response at the immediate post-receptor level (right, lower). In cells without GIV (left), or when GIV levels are reduced after prolonged LPS stimulation, TLR4-TIR modules readily homodimerize, even at low concentrations of LPS stimulation. Consequently, macrophages mount pro-inflammatory signals (NFkB, CREB and MAPK; left, lower) and are sensitive to stimuli.

## Conclusion

To ensure immunity, but avoid diseases and limit pathology, inflammatory responses must be finetuned through continuous feedback that confine responses to the proverbial ‘Goldilocks zone’ (47). The major discovery we report here is how macrophages confine TLR4 signaling and prevent an overzealous inflammatory response using a multi-modular scaffold, GIV. Prior work has eluded that balanced immune responses are brought about when the immune system dynamically responds to cues from both host and pathogen across multiple scales (47). Our findings showing that binding of GIV to TIR modules, thereby impacting multi-TLR signaling, could represent one such example of a multi-scale point of convergence for cues from host (levels of expression and immunomodulatory actions of GIV) and from diverse pathogens (ligands of multiple TLRs). GIV’s ability to bind some TIR modules (TLR4, TRAM, MAL) but not others (other TLRs, MyD88) *via* a single conserved SLIM suggests that GIV may function as a signal convergence/integration point for TLR crosstalk during pathogen-sensing. Whether other immune sensors and/or receptors (i.e., cytokine receptors, Nucleotide-binding oligomerization domain (**NOD**)-domain containing receptors, etc) also converge similarly, and if so, whether such convergence requires the same SLIM remains to be studied.

By elucidating the role of GIV, a non-canonical GEF for Gαi proteins, our findings also provide insights into a hitherto unknown mechanism of signal integration and convergence between TLRs and heterotrimeric G proteins. The role of the G-protein coupled receptor (GPCR)→G protein→cAMP signaling cascade in macrophage differentiation and in shaping macrophage responses to pathogens and injury-related danger molecules is well recognized (48). It has also been widely accepted that cross-talk between the GPCR→cAMP pathway and the TLRs impact LPS-triggered inflammatory responses (28, 49–51). By demonstrating that GIV modulates the Gαi→cAMP→CREB cascade during LPS/TLR4 signaling, we define a GPCR-independent regulation of cAMP in macrophages that may serve as a non-canonical mechanism for crosstalk between the LPS/TLR and G protein/cAMP pathways. Our finding that cAMP is increased in the hyper-reactive GIV-depleted macrophages is consistent with GIV’s previously described biochemical function (i.e., a GEF for Gαi), and with studies that have shown elevated levels of cAMP in response to LPS are responsible for the rapid induction of IL-6 production (28, 29). However, the impact of enhanced cAMP and CREB phosphorylation in GIV-depleted macrophages on downstream signaling and transcriptional responses remains unresolved. We propose that GIV-dependent suppression of cAMP in response to LPS may represent a non-canonical, anti-inflammatory cascade in macrophages responding to infectious stimuli (see legend, **Fig 8**).

Besides revealing the mechanisms underlying GIV’s immunomodulatory role, we have also demonstrated that GIV-derived peptides may serve as effective tools for mimicking such immunomodulatory role. In doing so, we have provided proof-of-concept that mechanistic insights gained herein can be exploited for the development of TLR4-inhibitory peptides, which could inspire immunomodulatory peptide-mimetic therapies.

Overall, this work ushers in new insights into how macrophage inflammatory responses are finetuned by the nature of protein complexes at the immediate-post receptor level and opens the door for the development of a new interface to target for immunomodulatory therapies.

## Materials and Methods

Methods related to Plasmid constructs, Protein expression and purification, Cell culture and immunoblotting, Generation of stable cell lines, Generation of conditional GIV KO mouse lines, *In vitro* Pulldown and Co-immunoprecipitation (Co-IP), RNA isolation, qPCR and RNA Sequencing, Cyclic AMP and NFkB measurements, Alignment of TIR domain sequences, Statistical Analysis and Replications are detailed in SI Appendix.

### Lipopolysaccharide (LPS) stimulation and Bacterial infection

For LPS stimulation experiments, cells were seeded (12-well plate: 2.5×10^5^ cells, 6-well plate: 5×10^5^ cells) and incubated overnight at 37°C before stimulation with LPS (dose and stimulation times indicated in figures and legends). For live microbe infection experiments, bacteria were maintained and cultured in accordance with ATCC protocols (52). Cells were infected at a multiplicity of infection (moi) of 1 for *E. coli* and 10 for *Salmonella*. For RNA readouts, cells were washed once with 1X PBS and stored a −80°C in Trizol. For supernatant cytokine analysis (ELISA), absolute levels of IL-6, IL-10, and TNFα were quantified using ELISA MAX or OptEIA ELISA kits (details in table).

### Bacterial infection of mice

Bacterial sepsis in mice was induced by injection of *E. coli* K1 strain RS218. For survival experiments, 9-week-old female Girdin floxed x LysMcre and littermate control WT mice were injected I.P. with 1×10^8^ colony forming units (cfu) *E. coli* and mouse survival was recorded for 24hrs following injection. For measurement of serum IL-6, IL-10 and IL-1β levels, serum was collected 3 hr after injection and cytokines were quantified by ELISA (R&D systems) following the manufacturer’s protocol.

### DSS colitis

Seven to eight-week-old GIV fl/fl x LysMcre or GIV fl/fl littermate controls were given either normal drinking water or 2% dextran sodium sulfate (DSS) for 7 days, followed by 7 days recovery with normal drinking water. DAI was calculated as done previously (53). Histology scoring was carried out as previously (52).

### Computational modeling of GIV-TLR interactions

Models were built by homology using the TIR-domain structures of TLR1 (54), TLR6 (55), and TLR10 (56) as templates. Homology modeling was performed in ICM (57, 58). The position of the conserved Pro-Gly motif of the GIV(1749-1761) peptide was inherited from the corresponding BB-loop motif in the homotypic TIR-domain homodimer of TLR10 (55) and the heterotypic TIR-domain homodimer of MAL/TIRAP (40). The peptide was built *ab initio*, tethered to the respective Pro-Gly positions, and its conformations were extensively sampled (> 10^8^ steps) by biased probability Monte Carlo (BPMC) sampling in internal coordinates, with the TLR4 TIR domain represented as a set of energy potentials precalculated on a 0.5 Å 3D grid and including Van der Waals potential, electrostatic potential, hydrogen bonding potential, and surface energy. Following such grid-based docking, the peptide poses were merged with full-atom models of the TLR4 TIR domain, and further sampling was conducted for the peptide and surrounding side chains of the TLR4 residues.

## Acknowledgments

This work was supported by National Institutes for Health (NIH) grants AI141630, CA100768 and CA160911 (to P.G), DK107585, CTSA/NCATS grant UL1TR001442 pilot award program and DiaComp Pilot and Feasibility award (to SD). PG and SD were also supported by the Helmsley Charitable Trust and Foundation for the Crohn’s disease project grant. L.S was supported by National Institutes for Health (NIH) training grant (DK 0070202). We would like to thank Dr. Christopher Glass (UCSD) and Dr. Gordon Gill (UCSD) for their critical feedback during the preparation of this manuscript.

## Contributions

P.G and L.S contributed to study conception and design. L.S, G.D.K, J.T., Y.C, J.C., V.C., and J.O., acquired, analyzed, and interpreted data. R.P. performed bioinformatical analyses. I.K. performed, analyzed, and interpreted computational homology modeling experiments. P.G and S.D. supervised work. S.D. and V.N. provided key reagents and protocols. P.G. and L.S. prepared data figures and wrote the manuscript. P.G. acquired funding for these studies.

## Corresponding author

Correspondence to Pradipta Ghosh

