## Supplementary Online Materials for "TLR4 signaling and macrophage inflammatory responses are dampened by GIV/Girdin"

### Appendix

#### Materials and Methods

##### Key Resource Table

| REAGENT or RESOURCE | SOURCE | IDENTIFIER |
|---|---|---|
| **Antibodies** | | |
| Rabbit polyclonal anti-GIV CT (1:500) | Santa Cruz Biotechnology | N/A |
| Rabbit polyclonal anti-Gαi3 (C-10) (1:500) | Santa Cruz Biotechnology | N/A |
| Rabbit polyclonal anti-β-tubulin (1:1000) | Santa Cruz Biotechnology | sc-9104 |
| Mouse monoclonal anti-TLR4 (1:500) | Santa Cruz Biotechnology | sc-293072 |
| Mouse monoclonal anti-actin (C2) (1:500) | Santa Cruz Biotechnology | sc-8432 |
| Rabbit polyclonal anti-NFκB (C-20) (1:500) | Santa Cruz Biotechnology | sc-372 |
| Normal mouse IgG | Santa Cruz Biotechnology | sc-2025 |
| Mouse monoclonal anti-poly-HIS (1:500) | Sigma-Aldrich | H1029 |
| Rabbit polyclonal anti-MyD88 (1:500) | Proteintech | 23230 |
| Rabbit polyclonal anti-Girdin coiled-coil (1:500) | Millipore-Sigma | ABT80 |
| Rabbit monoclonal anti-Phospho-NFκB p65 (Ser536) (93H1) (1:1000) | Cell Signaling Technology | #3033 |
| Rabbit monoclonal anti-Phospho-CREB (Ser133) (87G3) (1:1000) | Cell Signaling Technology | #9198 |
| Mouse monoclonal anti-CREB (86B10) (1:1000) | Cell Signaling Technology | #9104 |
| Rabbit monoclonal anti-Phospho-AKT (Ser473) (D9E) XP® (1:1000) | Cell Signaling Technology | #4060 |
| Mouse monoclonal anti-AKT (pan) (40D4) (1:1000) | Cell Signaling Technology | #2920 |
| Rabbit anti-Phospho-p44/42 MAPK (Erk1/2) (Thr202/Tyr204) (1:1000) | Cell Signaling Technology | #9101 |
| Mouse monoclonal anti-p44/42 MAPK (Erk1/2) (L34F12) (1:1000) | Cell Signaling Technology | #4696 |
| IRDye 800CW Goat anti-Mouse IgG Secondary (1:10,000) | LI-COR Biosciences | 926-32210 |
| IRDye 680RD Goat anti-Rabbit IgG Secondary (1:10,000) | LI-COR Biosciences | 926-68071 |
| **Biological Samples and Cell Lines** | | |
| RAW 264.7 | ATCC | TIB-71 |
| LysMcre mice (B6.129P2-Lyz2$^{tm1(cre)Ifo}$/j) | The Jackson Laboratory | 004781 |
| Girdin flox mice | *Asai et al. Biochem. Biophys. Res. Commun.*, 2012 | N/A |
| Escherichia coli K12 strain DH10B, pTransSacB | ATCC | PTA-5105 |
| *Salmonella enteric* serovar Typhimurium strain SL1344 | ATCC | 700720 |
| **Chemicals and Reagents** | | |
| Puromycin | Life Technologies | A1113803 |
| Polyethylenimine (PEI) | Polysciences, Inc | #23966 |
| TransIT®-LT1 Transfection Reagent | Mirus | 2300 |
| GIV TIR-peptide (RDVLPGT) | Biopeptide | N/A |
| Scrambled TIR-peptide (PTDLVRG) | Biopeptide | N/A |

| | | |
|---|---|---|
| Cell penetrating GIV TILL-peptide (RQIKIWFQNRRMKWKKKPEFLRPGPRKT) | Lifetein | N/A |
| Cell penetrating scrambled TILL-peptide (RQIKIWFQNRRMKWKKLPERKRPTKGPF) | Lifetein | N/A |
| Lipopolysaccharide (E. coli O111:B4) | Sigma-Aldrich | L4391 |
| Mouse TLR1-9 Agonist Kit | Invivogen | #tlrl-kit1mw |
| cAMP-Screen™ Cyclic AMP Immunoassay System | Applied Biosystems | 4412182 |
| Duel-luciferase® Reporter Assay System | Promega | E1910 |
| HisPur™ Cobalt Resin | Thermo Scientific | 89964 |
| Glutathione Sepharose® 4B | Sigma-Aldrich | GE17-0756-04 |
| Protein A Agarose | ThermoFisher | 15918014 |
| Protease inhibitor cocktail | Roche | 11 873 580 001 |
| Tyr phosphatase inhibitor cocktail | Sigma-Aldrich | P5726 |
| Ser/Thr phosphatase inhibitor cocktail | Sigma-Aldrich | P0044 |
| PVDF Transfer Membrane, 0.45μM | Thermo Scientific | 88518 |
| PowerUp™ SYBR™ Green Master Mix | Applied Biosciences | A25741 |
| qScript™ cDNA SuperMix | QuantaBio | 101414 |
| Direct-zol RNA Miniprep Kit | Zymo Research | R1051 |
| TRIzol™ Reagent | Invitrogen | 15596018 |
| ELISA MAX™ Deluxe Set Mouse IL-6 | BioLegend | 431304 |
| ELISA MAX™ Deluxe Set Mouse IL-1β | BioLegend | 432604 |
| ELISA MAX™ Deluxe Set Mouse IL-10 | BioLegend | 431414 |
| OptEIA™Mouse TNF (Mono/Mono) ELISA Set | BD Biosciences | 555268 |
| Dextran Sulfate Sodium Salt (Colitis Grade) | MP Biomedicals, LLC | 160110 |
| Hemoccult II | Beckman Coulter | 61130 |
| Zinc Formalin Fixative | Sigma-Aldrich | Z2902 |
| **Plasmids** | | |
| pET-28b-GIV-CT-WT (aa 1623-1870) | *Garcia-Marcos et al.,* 2009 | N/A |
| pET-28b-GIV-CT-WT (aa 1660-1870) | *Garcia-Marcos et al.,* 2009 | N/A |
| pET-28b-GIV-CT-WT (aa 1790-1870) | *Garcia-Marcos et al.,* 2009 | N/A |
| pET-28b-GIV-CT-K1749A (aa 1660-1870) | This paper | N/A |
| pET-28b-GIV-CT-EFL1751-1753AAA (aa 1660-1870) | This paper | N/A |
| pET-28b-GIV-CT-PG1755-1756AA (aa 1660-1870) | This paper | N/A |
| pET-28b-GIV-CT-K1749A (aa 1660-1870) | This paper | N/A |
| pET-28b-GIV-CT-PTB (PGxA) (aa 1660-1870) | This paper | N/A |
| pET-28b-Gαi3 (aa) | *Ghosh et al., 2009 PNAS* | N/A |
| pGEX-4T1-TLR4 (aa 676-835) | *Park et al., 2004* | N/A |
| pGEX-4T1-TLR4-Q683A (aa 676-835) | This paper | N/A |
| pGEX-4T1-TLR4-Q683R (aa 676-835) | This paper | N/A |
| pGEX-4T1-TLR4-E685A (aa 676-835) | This paper | N/A |
| pGEX-4T1-TLR4-E683R (aa 676-835) | This paper | N/A |
| pGEX-4T1-TLR4-Y709A (aa 676-835) | This paper | N/A |
| pGEX-4T1-TLR4-Y709F (aa 676-835) | This paper | N/A |
| pGEX-4T1-TLR4-Y709E (aa 676-835) | This paper | N/A |
| pGEX5-MyD88-TIR (aa163-296) | *Carlsson et al,. 2016* | N/A |
| pGEX-4T2-TRAM-TIR | *Lysakova-Devine et al,.* 2010 | N/A |
| pGEX-4T2-MAL-TIR | *Lysakova-Devine et al,.* 2010 | N/A |

| | | |
|---|---|---|
| pGL3basic- kappaB-luciferase (NFKB reporter) | *Lysakova-Devine et al,.* 2010 | N/A |
| pGEX6p1-TLR2 (aa641-784) | This paper | N/A |
| pGEX6p1-TLR1 (aa635-779) | This paper | N/A |
| pGEX6p1-TLR6 (aa640-784) | This paper | N/A |
| pET-28b-TLR4-TIR (aa676-835) | This paper | N/A |
| pLKO.1-GIV shRNA 1 (AAGAAGGCTTAGGCAGGAATT) | *Bhandari et al,.*2015 | N/A |
| pLKO.1-GIV shRNA 2 (GAAGGAGAGGCAACTGGAT) | *Bhandari et al,.*2015 | N/A |
| pLKO.1-scrambled shRNA (GGATTGAGATCAGAAGATAGC) | *Bhandari et al,.*2015 | N/A |
| **Primers** | Forward primer (5'→ 3') | Reverse primer (3'→ 5') |
| Girdin floxed screening primers | TGTCAGTTGTCAACTCTG AGC | ACACTGGTTGCTCTTCCA CAGTACTCTG |
| LysMcre screening primers | WT: TTACAGTCGGCCAGGCT GAC Mut: CCCAGAAATGCCAGATTA CG | CTTGGGCTGCCAGAATTT CTC |
| Mouse IL-6 qPCR primers | TGGAGTCACAGAAGGAG TGGCTAAG | TCTGACCACAGTGAGGAA TGTCCAC |
| Mouse IL-1β qPCR primers | GCCTTGGGCCTCAAAGG AAAGAATC | GGAAGACACAGATTCCAT GGTGAAG |
| Mouse TNFα qPCR primers | ATAGCTCCCAGAAAAGCA AGC | CACCCCGAAGTTCAGTAG ACA |
| Mouse IL-10 qPCR primers | CCCTGGGTGAGAAGCTG AAG | CACTGCCTTGCTCTTATT TTCACA |
| Mouse IFNβ qPCR primers | AAGAGTTACACTGCCTTT GCCATC | CACTGTCTGCTGGTGGA GTTCATC |
| Mouse 18S qPCR primers | GTAACCCGTTGAACCCCA TT | CCATCCAATCGGTAGTAG CG |
| **Software** | | |
| Prism | GraphPad | https://www.graphpad.com/scientific-software/prism/ |
| Molsoft | Molsoft, LLC | https://www.molsoft.com/index.html |
| Illustrator | Adobe | https://www.adobe.com/products/illustrator.html |
| ImageStudio Lite | LI-COR | https://www.licor.com/bio/image-studio-lite/ |

**Plasmid constructs**

Cloning of GIV-CT (aa 1623-1870, 1660-1870, and 1790-1870) into pET28b (His-GIV CT) were previously described (59). His-GIV-CT mutants (detailed in table) were generated by site-directed mutagenesis using QuickChange kit (Stratagene) and specific primers (sequence available in Key Resource Table) as per the manufacturer's protocols. Cloning of Gαi3 into pET28b was previously described (60). GST-tagged TLR4 was obtained from Yun Soo Bae (61). GST-tagged MyD88-TIR was obtained from Bernadette Byrne (62). GST-tagged TRAM and MAL, and NFkB reporter plasmids were obtained from Andrew Bowie (63). GST-tagged TLR1, TLR2, and TLR6 were cloned by amplifying TIR regions from THP-1 macrophage (human) cDNA and subcloned into pGEX-6p1 using BamHI and EcoRI restriction sites. His-tagged TLR4 was generated by subcloning human TLR4 (from GST-TLR4 construct described above) into pET28b using BamHI and EcoRI restriction sites. Cloning of GIV shRNA constructs was previously described (64).

**Protein expression and purification**

GST and His-tagged proteins were expressed in E. coli strain BL21 (DE3) and purified as previously described (59, 60). Briefly, cultures were induced using 1mM IPTG overnight at 25°C. Cells were then pelleted and resuspended in either GST lysis buffer (25 mM Tris-HCL (pH 7.4), 20 mM NaCl, 1 mM EDTA, 20% (vol/vol) glycerol, 1% (vol/vol) Triton X-100, protease inhibitor cocktail) or His lysis buffer (50 mM $NaH_2PO_4$ (pH7.4), 300 mM NaCl, 10 mM imidazole, 1% (vol/vol) Triton-X-100, protease inhibitor cocktail). Cells were lysed by sonication, and lysates were cleared by centrifugation at 12,000 x g at 4°C for 30 mins. Supernatant was then affinity purified using glutathione-Sepharose 4B beads or HisPur Cobalt Resin, followed by elution, overnight dialysis in PBS, and then storage at -80°C.

**Cell culture, transfection, lysis, and immunoblotting**

The RAW 264.7 cell line was obtained from and cultured according to American Type Culture Collection (ATCC) guidelines. Transfection, lysis, and immunoblotting were carried out as described previously (65). Whole-cell lysates were prepared after washing cells with cold PBS before resuspending and boiling them in sample buffer. For immunoblotting, protein samples were separated by SDS-PAGE and transferred to polyvinylidene fluoride membranes. Membranes were blocked with 5% non-fat milk or 5% BSA (when probing for phosphorylated proteins) in PBS. The membrane was stained with Ponceau S to visualize bait proteins, washed, blocked, and incubated with primary antibody solutions overnight at 4°C (dilutions for each primary antibody detailed in table). Washed blots were then incubated with infrared secondary antibodies (detailed in table) for 1 hr at room temperature. Infrared imaging with two-color detection and quantification were performed using a Li-Cor Odyssey imaging system and analysis was performed with Image StudioLite software.

**Generation of stable cell lines**

ShRNA control and shRNA GIV RAW 264.7 stable cells lines were generated by lentiviral transduction followed by selection with puromycin as described previously (66). Lentiviral packaging was performed in HEK293T cell by co-transfecting shRNA constructs with psPAX2 and pMD2G plasmids (4:3:1 ratio) using Mirus LT1. The medium was changed after 24 hr, and virus-containing media was collected after 36-48 hr, centrifuged, and filtered through a 0.45 μM filter. Fresh virus-containing media was diluted 1:4 with RAW 264.7 media (DMEM, 10% FBS) and polybrene (6 μg/ml final concentration) was added. Lentiviral mixture was added to RAW 264.7 macrophages ($2.5x10^5$ cells seeded in 6-well plate), spun at 800 x g at room temperature for 30 min, and transferred to cell culture incubator and media was changed after 4 hrs. Puromycin (2.5 μg/ml) was added 48 hrs post-transduction for selection. Depletion of endogenous GIV was confirmed by immunoblotting with GIV-CC (ABT80) rabbit antibody.

**Generation of conditional GIV KO mouse lines**

All breeding and mouse experimentation were done in accordance with the rules and regulations of the Institutional Animal Care and Use Committee (IACUC), at the University of California, San Diego and all measures were taken to use animal subjects efficiently and humanely. Girdin floxed mice were a generous gift from Dr. Masahide Takahashi (Nagoya University, Japan) and were described previously (11). LysMcre mice (B6.129P2-Lyz2$^{tm1(cre)Ifo}$/j) were purchased from The Jackson Laboratory. Girdin floxed x LysMcre mice were generated by us and were maintained as homozygous floxed (fl/fl) and heterozygous LysMcre. Mice were genotyped by PCR (primers in table).

**RNA isolation, quantitative PCR, and RNA sequencing**

All RNA was isolated using Direct-zol RNA Miniprep Kit using manufactures protocol from samples collected in Trizol reagent. RNA concentration and purity were quantified using a Nanodrop Microvolume Spectrophotometer. 500 ng RNA was used for RT-PCR using qScript cDNA SuperMix kit and manufacturers protocol. cDNA was diluted 1:5 with dd$H_2O$ and qPCR was carried out using 2X PowerUp SYBR Green Master Mix (primer sequences listed in table). The cycle threshold (Ct) of target genes was normalized to 18S housekeeping gene, relative expression of mRNA was calculated using the DDCt method, and results expressed as fold-change. For determining which genes were differentially expressed by RNAseq, transcript-level abundance of paired-end RNA-seq data was estimated by Salmon (1.1.0) using the mouse transcriptome from Genecode (vM24). Transcript-level quantification was aggregated to the gene-level with Tximport (1.14.2). The resulting gene counts were used as an input to DESeq2 (1.26.0). Differentially expressed genes below a BH adjusted p-value of 0.05 were considered significant. Volcano plots were created using the Enhanced Volcano R package (1.4.0). Gene symbols were added by BiomaRt (2.42.1). Snakemake (5.10.0) was used as a workflow management system for processing fastq files with salmon.

**Lipopolysaccharide (LPS) stimulation and Bacterial infection**

For LPS stimulation experiments, cells were seeded (12-well plate: $2.5x10^5$ cells, 6-well plate: $5x10^5$ cells) and incubated overnight at 37°C before stimulation with LPS (dose and stimulation times indicated in figures and legends). For live microbe infection experiments (*E. coli* and *Salmonella*), bacteria were maintained and cultured in accordance with ATCC protocols as was previously described (52). For bacterial culture, a single colony was inoculated into LB broth and grown for 8 hr under aerobic conditions in an orbital shaking incubator at 150 rpm and then under oxygen-limiting conditions overnight. Under these conditions, bacteria correspond to $5\text{-}7x10^8$ colony forming units (CFU), where OD 0.5 is equivalent to $5x10^8$. Cells were infected at a multiplicity of infection (moi) of 1 for *E. coli* and 10 for *Salmonella*. For RNA readouts, cells were washed once with 1X PBS and 0.5ml Trizol was added directly to the well before collection and storage at -80°C. For supernatant cytokine analysis (ELISA), supernatant was collected at indicated times, centrifuged at 13,000 x g for 10 min, and absolute levels of IL6, IL10, and TNFα were quantified using ELISA MAX or OptEIA ELISA kits (details in table) using manufacturers protocols. For cell lysate analysis (Western blot), cells were washed 1X with PBS, scraped from well, and processed as described above.

**Bacterial infection of mice**

Bacterial sepsis in mice was induced by injection of *E. coli* K1 strain RS218. The *E. coli* culture was grown overnight in Luria broth (LB) medium at 37°C with shaking. The bacterial culture was diluted 1:50 in fresh LB, grown to mid-log phase, washed twice with PBS and reconstituted in PBS to yield the appropriate inoculum. For survival experiments, 9-week-old female Girdin floxed x LysMcre and littermate control WT mice were injected with $1x10^8$ colony forming units (cfu) *E. coli* and mouse survival was recorded for 24hrs following injection. For measurement of serum IL-6, IL-10 and IL-1β levels, mice were injected with $1x10^8$ colony forming units (cfu) *E. coli* and at 3 hr after injection 80 µl of blood was collected by submandibular bleeding using a lancet into a serum separating blood collection tubes (BD) that were spun at 1500 × *g* for 10 min to separate serum. Serum cytokines were quantified by specific ELISA (R&D systems) following the manufacturer's protocol.

**DSS colitis**

Seven to eight-week-old GIV fl/fl x LysMcre or GIV fl/fl littermate controls were given either normal drinking water or 2% dextran sodium sulfate (DSS) for 7 days, followed by 7 days recovery with normal drinking water. Water levels were monitored to ensure equal volumes of water were consumed between treatment groups. Weight was monitored daily. Disease activity index (DAI) was calculated using by scoring stool consistency (0-4), rectal bleeding (0-4), and weight loss (0-4) as previously published (53). Mice were sacrificed on the day 14, and colon length was measured. Colon samples were prepared as Swiss-rolls, fixed in formalin, embedded in paraffin, and cut into sections. Sections were stained with hematoxylin and eosin and evaluated for mononuclear infiltrates, submucosal edema, surface erosions, inflammatory exudates, and presence of crypt abscesses and scored as done previously (52).

**Measurement of cellular cAMP levels**

RAW 264.7 cells ($2.5x10^5$ cells/well in 12-well plate) were incubated with 200 μM isobutyl methyl xanthine (IBMX) for 20 min, followed by 1 hr LPS (100 ng/ml) stimulation. Cells were lysed and cAMP levels assessed by cAMP-Screen Cyclic AMP Immunoassay System using manufacturers protocols, and data expressed as pmol cAMP/ml.

**NFkB reporter assay**

RAW 264.7 cells ($5x10^4$ cells/well in 96-well plate) were transfected with 100 ng NFkB reporter plasmid and 20 ng Renilla luciferase control plasmid. 24 hr after transfection, cells were stimulated with LPS (100 ng/ml) for 6hr and NFkB activity was assessed using the Dual-luciferase Reporter Assay System using manufacturers protocol.

##### *In vitro* Pulldown and Co-immunoprecipitation (Co-IP)

For *in vitro* pulldown assays, purified GST-tagged proteins from E. coli were immobilized onto glutathione-Sepharose beads by incubating with binding buffer (50 mM Tris-HCl (pH 7.4), 100 mM NaCl, 0.4% (vol/vol) Nonidet P-40, 10 mM MgCl2, 5 mM EDTA, 2 mM DTT) for 60 min at 4°C. GST-protein bound beads were washed and incubated with purified His-tagged proteins resuspended in binding buffer for 4 hrs at 4°C. After binding, bound complexes were washed four times with 1 ml phosphate wash buffer (4.3 mM $Na_2HPO_4$, 1.4 mM $KH_2PO_4$(pH 7.4), 137 mM NaCl, 2.7 mM KCl, 0.1% (vol/vol) Tween-20, 10 mM $MgCl_2$, 5 mM EDTA, 2 mM DTT, 0.5 mM sodium orthovanadate) and eluted by boiling in Laemmli buffer (5% SDS, 156 mM Tris-Base, 25% glycerol, 0.025% bromophenol blue, 25% β-mercaptoethanol). For immunoprecipitation assays, cell were lysed in cell lysis buffer (20 mM HEPES (pH 7.2), 5 mM Mg-acetate, 125 mM K-acetate, 0.4% Triton X-100, 1 mM DTT, 0.5 mM sodium orthovanadate, Tyr phosphatase inhibitor cocktail, Ser/Thr phosphatase inhibitor cocktail, and protease inhibitor cocktail) using a 28G syringe, followed by centrifugation at 10,000 x g for 10 min. Cleared supernatants were then incubated with mouse anti-TLR4 or mouse IgG control antibodies at 4°C overnight. Protein A agarose beads were added to immuno-bound lysates and incubated for 1 hr at 4°C. Immunocomplex bound beads were washed with wash buffer (4.3 mM $Na_2HPO_4$, 1.4 mM $KH_2PO_4$(pH 7.4), 137 mM NaCl, 2.7 mM KCl, 0.1% (vol/vol) Tween-20, 10 mM $MgCl_2$, 5 mM EDTA, 2 mM DTT, 0.5 mM sodium orthovanadate) and eluted by boiling in Laemmli buffer (5% SDS, 156 mM Tris-Base, 25% glycerol, 0.025% bromophenol blue, 25% β-mercaptoethanol).

##### Alignment of TIR domain sequences

Amino acid sequences of TIR-domain containing proteins was retrieved from Ensembl Genome Browser (www.ensembl.org) and aligned using Clustal Omega Multiple Sequence Alignment tool (https://www.ebi.ac.uk/Tools/msa/clustalo/). Alignments were displayed using BoxShade Server (https://embnet.vital-it.ch/software/BOX_form.html).

##### Computational modeling of GIV-TLR interactions

Since the structure of the TLR4 TIR-domain is not available, models were built by homology using the TIR-domain structures of TLR1 (54), TLR6 (55), and TLR10 (56) as templates. Homology modeling was performed in ICM (57, 58). The position of the conserved Pro-Gly motif of the GIV(1749-1761) peptide was inherited from the corresponding BB-loop motif in the homotypic TIR-domain homodimer of TLR10 (55) and the heterotypic TIR-domain homodimer of MAL/TIRAP (40). The peptide was built *ab initio*, tethered to the respective Pro-Gly positions, and its conformations were extensively sampled (> $10^8$ steps) by biased probability Monte Carlo (BPMC) sampling in internal coordinates, with the TLR4 TIR domain represented as a set of energy potentials precalculated on a 0.5 Å 3D grid and including Van der Waals potential, electrostatic potential, hydrogen bonding potential, and surface energy. Following such grid-based docking, the peptide poses were merged with full-atom models of the TLR4 TIR domain, and further sampling was conducted for the peptide and surrounding side chains of the TLR4 residues.

**Statistical Analysis and Replications**

Statistical significance between datasets with three or more experimental groups was determined using one-way (or two-way in the case of DSS weight analysis) analysis of variance (ANOVA) including a Tukey's test for multiple comparisons. Statistical difference between two experimental groups was determined using a two-tailed unpaired t-test. For all tests, a p-value of 0.05 was used as the cutoff to determine significance. All experiments were repeated a least three times, and p-values are indicated in each figure. All statistical analysis was performed using GraphPad prism 8.

### Supplementary Figures

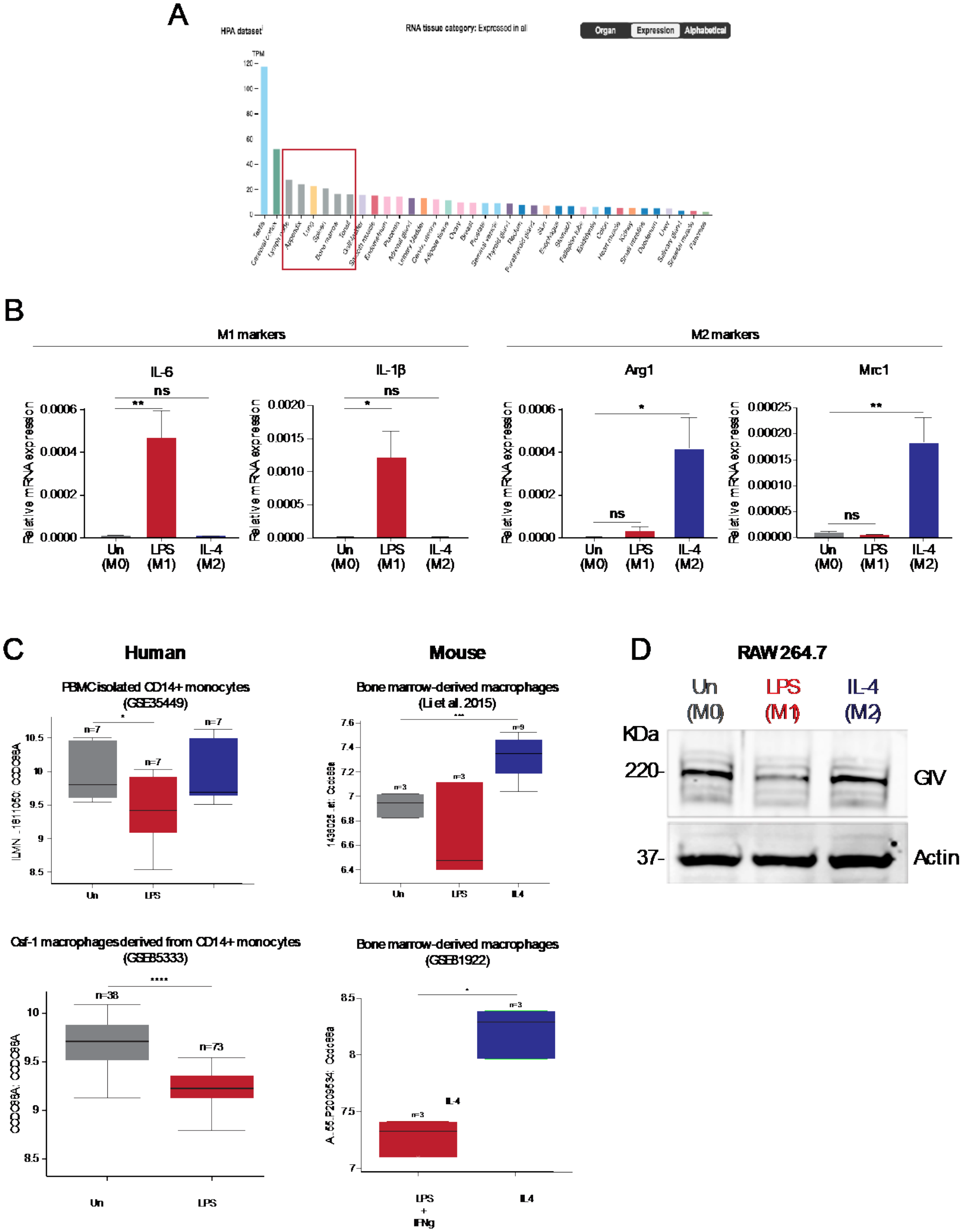

**Supplementary Figure 1: GIV expression in immune tissues and polarized macrophages. (A)** Bar graph, generated using Human Protein Atlas (www.proteinatlas.org), showing GIV transcript levels from various human tissues. Red box highlights GIV expression in immune tissues. **(B)** Immunoblot showing GIV levels in RAW 264.7 macrophages stimulated with either LPS (100ng/ml) or IL-4 (20ng/ml) for 24hrs. **(C)** Box plots of GIV transcript levels, curated from publicly available RNAseq datasets (Gene Expression Omnibus [GEO]), in human and mouse macrophages polarized under indicated pro- or anti-inflammatory conditions. **(D)** Bar graphs displaying transcript levels (qPCR) of M1 and M2 polarization markers in RAW 264.7 macrophages stimulation with LPS (100ng/ml) or IL-4 (20ng/ml) for 24hr. Results are from three independent experiments and displayed as mean ± S.E.M.. One-way ANOVA was used to determine significance. (*;p ≤ 0.05, **;p ≤ 0.01).

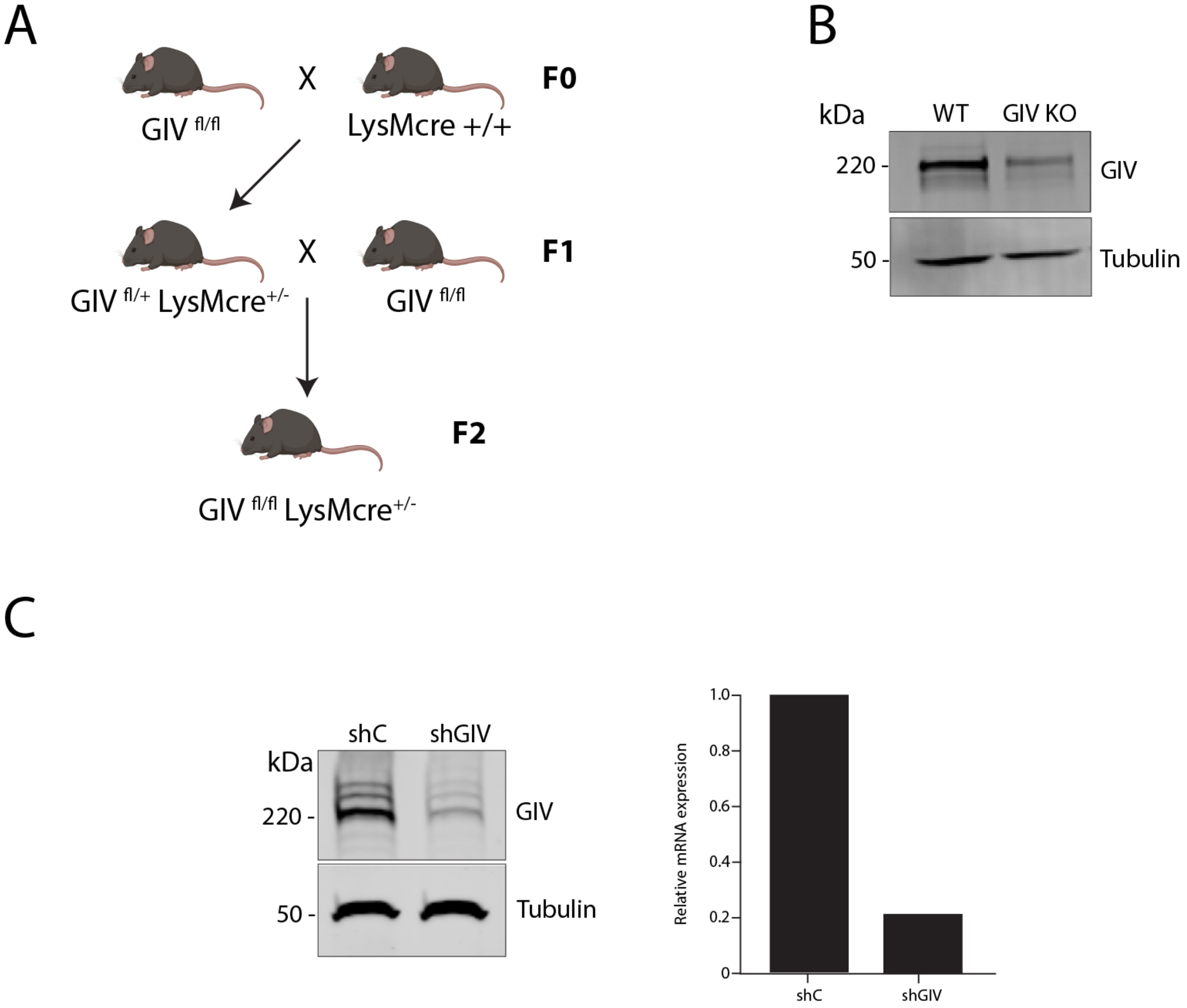

**Supplementary Figure 2: Validation of GIV-depletion model systems: (A)** Schematic of breeding scheme used to generate conditional myeloid-cell specific (LysMcre) GIV knockout mice. **(B)** Immunoblot confirming protein depletion in peritoneal macrophages isolated from GIV fl/fl x LysMcre mice. **(C)**. (left) Immunoblot confirming protein depletion in RAW 264.7 macrophage cell line using shRNA. (right) Bar graph of GIV mRNA levels in shRNA GIV-depleted macrophages relative to WT controls.

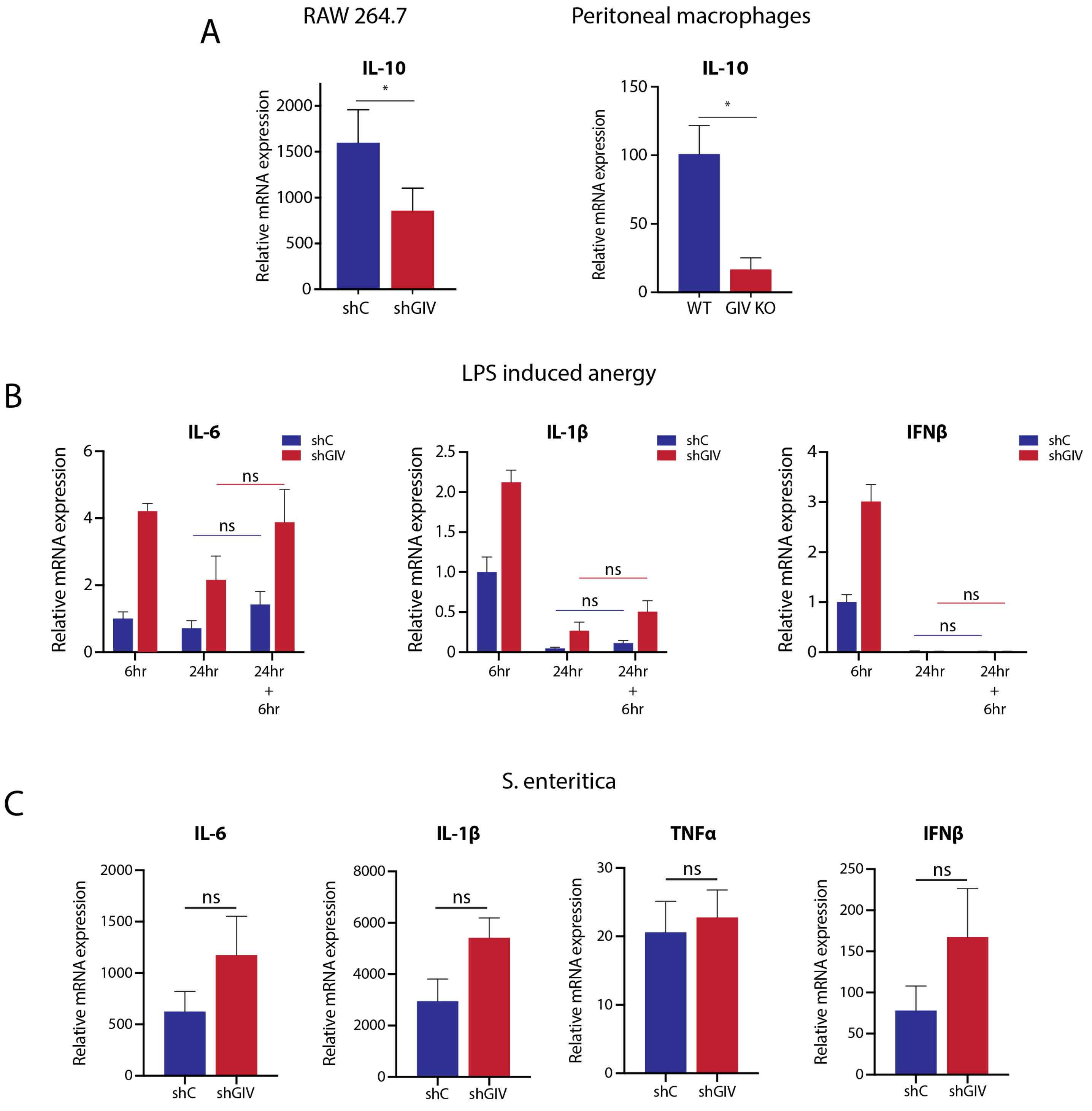

**Supplementary Figure 3: GIV depletion enhances pro-inflammatory cytokine response but is not required for LPS induced anergy. (A)** Bar graphs showing transcript levels of IL10 in GIV-depleted or control RAW 264.7 or peritoneal macrophages stimulated with LPS (100ng/ml, 6hr). **(B)** Bar graphs of pro-inflammatory cytokine transcript levels in RAW 264.7 macrophages either 1) simulated with LPS (100ng/ml) for 6hrs, 2) stimulated with LPS (100ng/ml) for 24hrs, or 3) stimulated with LPS (100ng/ml) for 24hr before re-challenge with LPS (100ng/ml) for 6hrs. **(C)** Bar graphs displaying cytokine transcript levels (qPCR) in GIV-depleted RAW 264.7 macrophages infected with live *Salmonella* (MOI=10) for 6hrs compared to controls. Results were from 3 independent experiments and displayed as mean ± S.E.M.

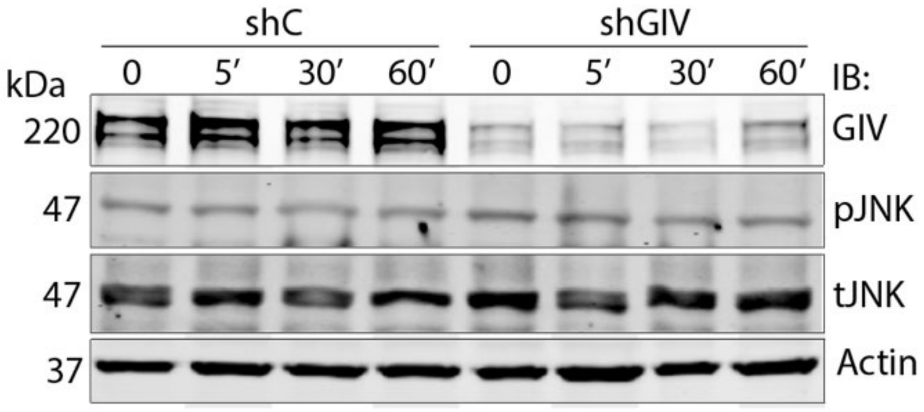

**Supplementary Figure 4: GIV depletion in macrophages does not impact JNK MAPK activation.** Immunoblot of whole-cell lysates from GIV-depleted or control RAW 264.7 macrophages stimulated with LPS (100 ng/ml) and probed for activation of indicated signaling pathways.

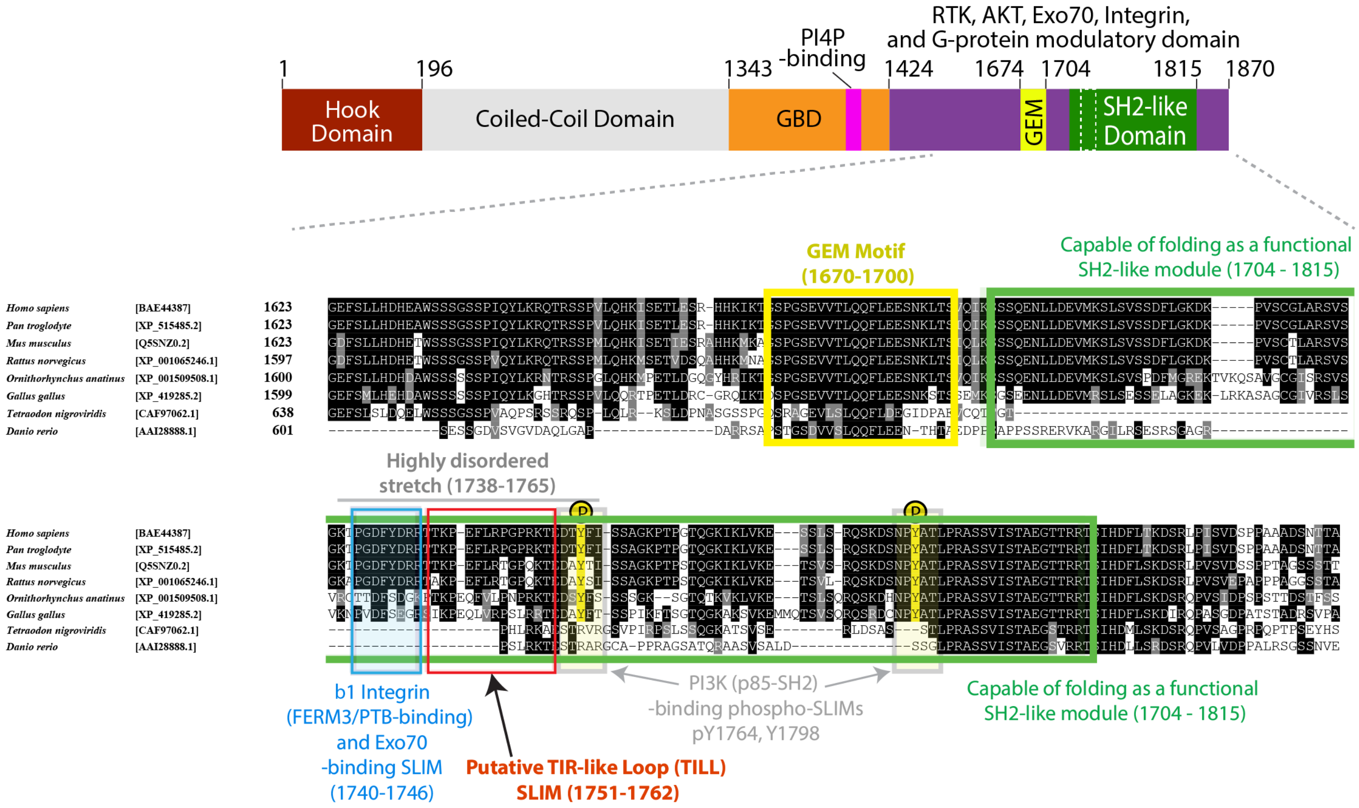

**Supplementary Figure 5: Short linear interaction motifs (SLIMs) within GIV's C-terminus. Top:** Bar diagram showing the various domains of GIV. GBD, G protein binding domain; GEM, Guanine nucleotide exchange modulator; SH2, Src-like homology; PI4P, phosphoinositol-4-phosphate. **Bottom**: Sequence of GIV's C-terminus showing all currently identified SLIMs. Putative TIR-like Loop (TILL) SLIM is highlighted in red.

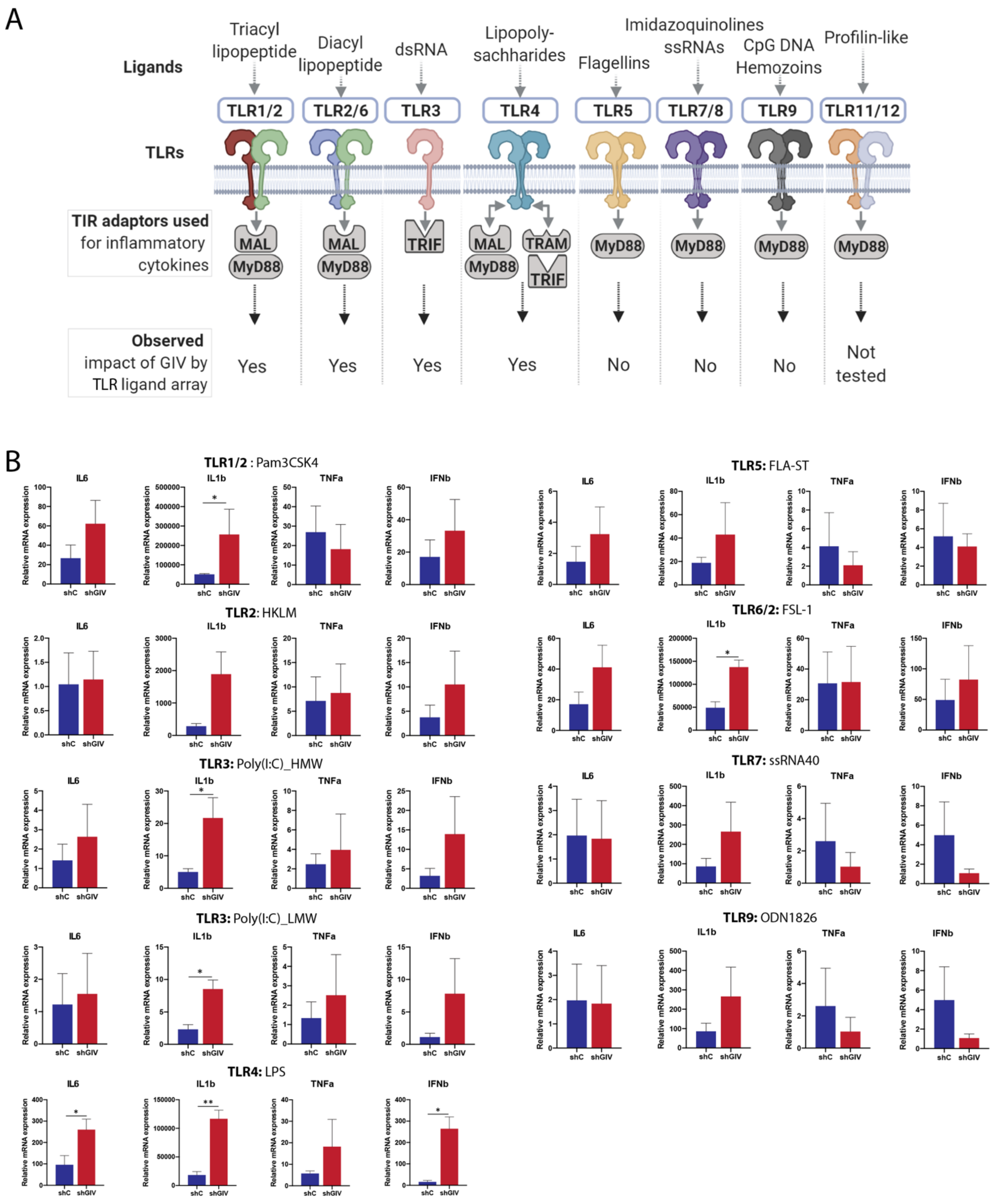

**Supplementary Figure 6: GIV impacts inflammatory responses downstream of TLRs that require the TIR-adaptors MAL, TRAM, and TRIF. (A)** Schematic summarizing results from TLR-array assay highlighting GIV's impact on TLR responses that require MAL, TRAM, and TRIF TIR adaptors. **(B)** Bar graphs showing transcript levels of pro-inflammatory cytokines in GIV-depleted or control RAW 264.7 macrophages in response to various TLR ligand stimulation for 6hr. Results are from 3 independent experiments and displayed as mean ± S.E.M. Students t-test was used to determine significance. (*;p≤0.05, **;p≤0.01, ***;p≤0.001, ****;p≤0.0001).

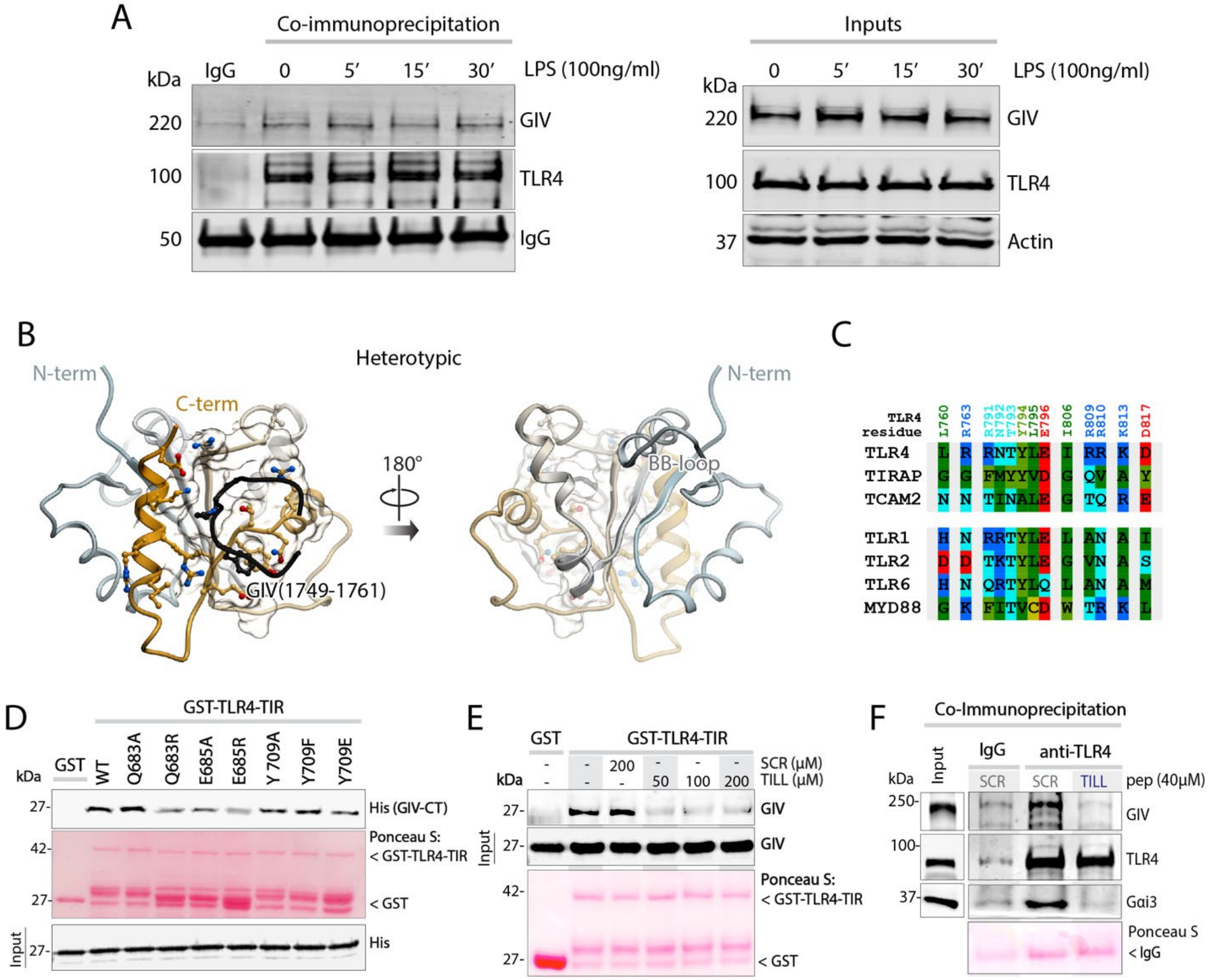

**Supplementary Figure 7: Heterotypic interaction model of GIV and TLR4-TIR. (A)** Endogenous TLR4 was immunoprecipitated from LPS stimulated (100ng/ml) RAW 264.7 lysates. Bound complex of TLR4 and GIV were visualized by immunoblot. Equal loading of IgG and anti-TLR4 were confirmed by immunoblot. **(B)** A model of hetrotypic interaction between GIV's TILL domain (black) and the C-terminal helix (orange) of TLR4's TIR-domain. **(C)** Sequence alignment TLR4 with other TLRs and TIR-adaptor proteins showing proposed contact residues between GIV and TLR4's C-terminal helix region. **(D)** Recombinant His-GIV-CT was used in a GST pulldown assay with either WT GST-TLR4-TIR or GIV•TLR4 interface mutants and bound GIV was visualized by immunoblot. **(E)** Recombinant His-GIV-CT was used in GST pulldown assay with GST or GST-TLR4-TIR and increasing amounts of TILL-peptide or scrambled (SCR) control peptide. Bound GIV was visualized by immunoblot. Equal loading of GST proteins was confirmed by Ponceau S staining. **(F)** Endogenous TLR4 was immunoprecipitated from RAW 264.7 lysates in the presence of either TILL-peptide or scrambled control. Bound complex of TLR4, GIV, and Gαi were visualized by immunoblot. Equal loading of IgG and anti-TLR4 were confirmed by Ponceau S staining.
